# Pre-existing resistant proviruses can compromise maintenance of remission by VRC01 in chronic HIV-1 infection

**DOI:** 10.1101/2020.02.14.940395

**Authors:** Ananya Saha, Narendra M. Dixit

## Abstract

Broadly neutralizing antibodies (bNAbs) of HIV-1 hold promise of eliciting long-term HIV-1 remission. Surprisingly, the bNAb VRC01, when administered concomitantly with the cessation of successful antiretroviral therapy (ART), failed rapidly in chronic HIV-1 patients. We hypothesized that the failure was due to VRC01-resistant strains that were formed before ART initiation, survived ART in latently infected cells, and were reactivated during VRC01 therapy. Current assay limitations preclude testing this hypothesis experimentally. We developed a mathematical model based on the hypothesis and challenged it with available clinical data. The model integrated within-host HIV-1 evolution, stochastic latency reactivation and viral dynamics with multiple dose VRC01 pharmacokinetics. With a virtual patient population, model predictions quantitatively captured data from two independent clinical trials. Accordingly, we attributed VRC01 failure to single-mutant VRC01-resistant proviruses in the latent reservoir triggering viral recrudescence, particularly during trough VRC01 levels. Accounting for pre-existing resistance may help bNAb therapies maximize HIV-1 remission.

## INTRODUCTION

Antiretroviral therapy (ART) for HIV-1 infection rapidly suppresses viremia to undetectable levels and curtails disease progression but is unable to eradicate the virus.^1^ Discontinuation of ART, even long after viremic control is established, typically leads to rapid viral rebound, often within days to weeks of discontinuation, and to progressive disease.^2^ The rebound is caused by a reservoir of latently infected cells^3^ that is formed soon after infection^4^. The reservoir is sustained by cell proliferation^5,6^, which can continue even when ART has stopped new infections, allowing the reservoir to exist long-term^7^. The reservoir can get reactivated stochastically and reignite infection once ART is stopped^8,9^. ART must therefore be administered lifelong. In a remarkable breakthrough, the VISCONTI study showed that when ART is administered early in infection, some individuals can maintain viremic control for many years after the cessation of ART^10^. This study has raised hopes of a functional cure, or long-term remission, of HIV-1, where the viremic control once established by ART can be maintained without lifelong treatment^11^. The success of ART in inducing post-treatment control, however, is small: only ~5-15% of the patients treated achieve lasting post-treatment control^12^. Enormous efforts are underway to improve this success rate^11^.

One strategy that holds promise is to administer broadly neutralizing antibodies (bNAbs) of HIV-1 for a short period post-ART, i.e., during an analytical treatment interruption (ATI)^13,14^. bNAbs target diverse viral genomic variants^15,16^, and are expected to suppress viremia arising from the reactivation of latently infected cells^17,18^. Simultaneously, they may engage the host immune system^19^, potentiating it to maintain the viremic control long-term^20,21^. Two recent clinical trials tested this strategy using the bNAb VRC01^17^. VRC01 targets the CD4 binding site on the HIV-1 envelope with high breadth and potency^15,22^. When it was administered to chronically infected patients who had achieved undetectable viremia with ART, the duration of viremic control was observed to increase only marginally, by a median of ~2-4 weeks, beyond historical controls^17^. In the historical control group, which was not treated post-ART, viral rebound occurred in ~2.6 weeks on average after ART interruption^23^. Why VRC01 was ineffective in maintaining remission longer is unclear. Unravelling the causes of this ineffectiveness is important to optimizing VRC01 usage, which is also in large clinical trials for preventing the transmission of infection^24^, and to expose potential vulnerabilities of other bNAbs.

VRC01 exhibits potent antiviral activity *in vivo*^25^, including at its trough levels in patients^26^. Yet, in the trials above^17^ viral rebound was observed in most patients when VRC01 levels in circulation were significantly higher than its *in vitro* suppressive concentration (or *IC*_50_), implicating the role of resistance. Mutations that confer resistance to bNAbs are well documented^27–30^. Indeed, VRC01-resistant strains were detected in the breakthrough viral populations in the trials above^17^. The rapid virological breakthrough during treatment, especially given the absence of circulating virions at the time of ART cessation and the potent activity of VRC01 against the wild-type, suggests that VRC01 resistance might have existed before VRC01 therapy. Here, we therefore hypothesized that pre-existing VRC01-resistant proviruses formed before ART and contained in the latent reservoir could underlie the failure of VRC01 therapy. The inability to detect minority viral variants as well as latently infected cells using current assays^31,32^ highlights the challenge associated with testing our hypothesis experimentally. Current assays can rarely detect variants below a frequency of ~1%^33^, implying that, given the prevalent estimates of the latent reservoir size of 10^5^ –10^8^ cells^4,34^, variants present in as many as 10^3^ latently infected cells may go undetected and be responsible for therapy failure. Indeed, in many individuals who failed rapidly in the trials above^17^, resistance could not be detected pre-treatment. As an alternative approach, therefore, we resorted to mathematical modeling.

A number of mathematical models have been developed in recent years to describe latent cell infection and dynamics and its role in the outcomes of treatments^35–46^. Models have also been constructed, independently, for describing viral evolution and drug resistance, especially in the context of ART^47–51^. Describing the failure of VRC01 therapy required integrating these two independent formalisms, of viral evolution and latent reservoir reactivation, a task not accomplished so far because of the complexity involved. HIV-1 evolution involves mutation, recombination, fitness selection, and random genetic drift, which together define the timing and speed of the development of drug resistance during ART^49–53^. Latent cell reactivation is an intrinsically stochastic process^8,43,44^, following which the virions released must establish lasting infection, which is not guaranteed^35,39,42^, especially in the presence of bNAbs. Here, we developed a framework that integrates these processes by recognizing that the dynamics of viral evolution and latent cell reactivation could be decoupled in the context of post-ART bNAb therapy. Viral evolution primarily occurs pre-ART, where viral and productively infected cell populations are large and the contribution from latently infected cells to the dynamics can be ignored. Latent cell reactivation leading to treatment failure occurs during bNAb therapy, when active viral replication is small and so viral evolution can be ignored. These processes are linked by VRC01 resistant strains, which are predominantly formed before therapy and are harbored in latently infected cells and could get reactivated during therapy. Developing a model with this strategy, we were able, for the first time, to capture data from human clinical trials involving bNAb-based interventions quantitatively, offering an explanation for the inadequate effectiveness of VRC01 in maintaining remission, and providing a framework for rational treatment optimization.

## RESULTS

### Mathematical model

We considered the scenario where chronically infected individuals with viremic control established with long-term ART are administered VRC01 during an ATI, as in recent clinical trials^17^. We developed a model to predict the ensuing remission times based on the hypothesis that viral strains resistant to VRC01 harbored in latently infected cells were responsible for virological breakthrough (Fig. 1). We provide an overview of the model here; details are in Methods.

We first considered a single infected individual (Fig. 1(a)). We estimated the diversity of the viral population in the individual at ART initiation using a detailed model of viral dynamics and evolution that considered target cells, free virions, and productively infected cells, and included mutation, cellular superinfection, recombination, and fitness selection based on the relative fitness of dominant VRC01 resistance mutations. From the diversity, we obtained the frequencies of productively infected cells containing different mutant proviruses resistant to VRC01. We let the latent reservoir harbor proviruses with the same frequencies, as has been done previously^53^. We recognized that of the latter cells, those harboring the most frequent mutant provirus were the most likely to re-establish infection. We employed a stochastic model to estimate the waiting time for the reactivation of such latent cells and tracked the ensuing dynamics during VRC01 therapy until the infection grew to detectable levels, at which point the therapy was deemed to have failed. During the simulations, we let the efficacy of VRC01 vary continuously with time based on its pharmacokinetic profile, which we estimated from independent fits to data. The parameter values employed for estimating the proviral frequencies and latent cell reactivation are listed in Tables 1 and 2, respectively.

**Figure 1:**
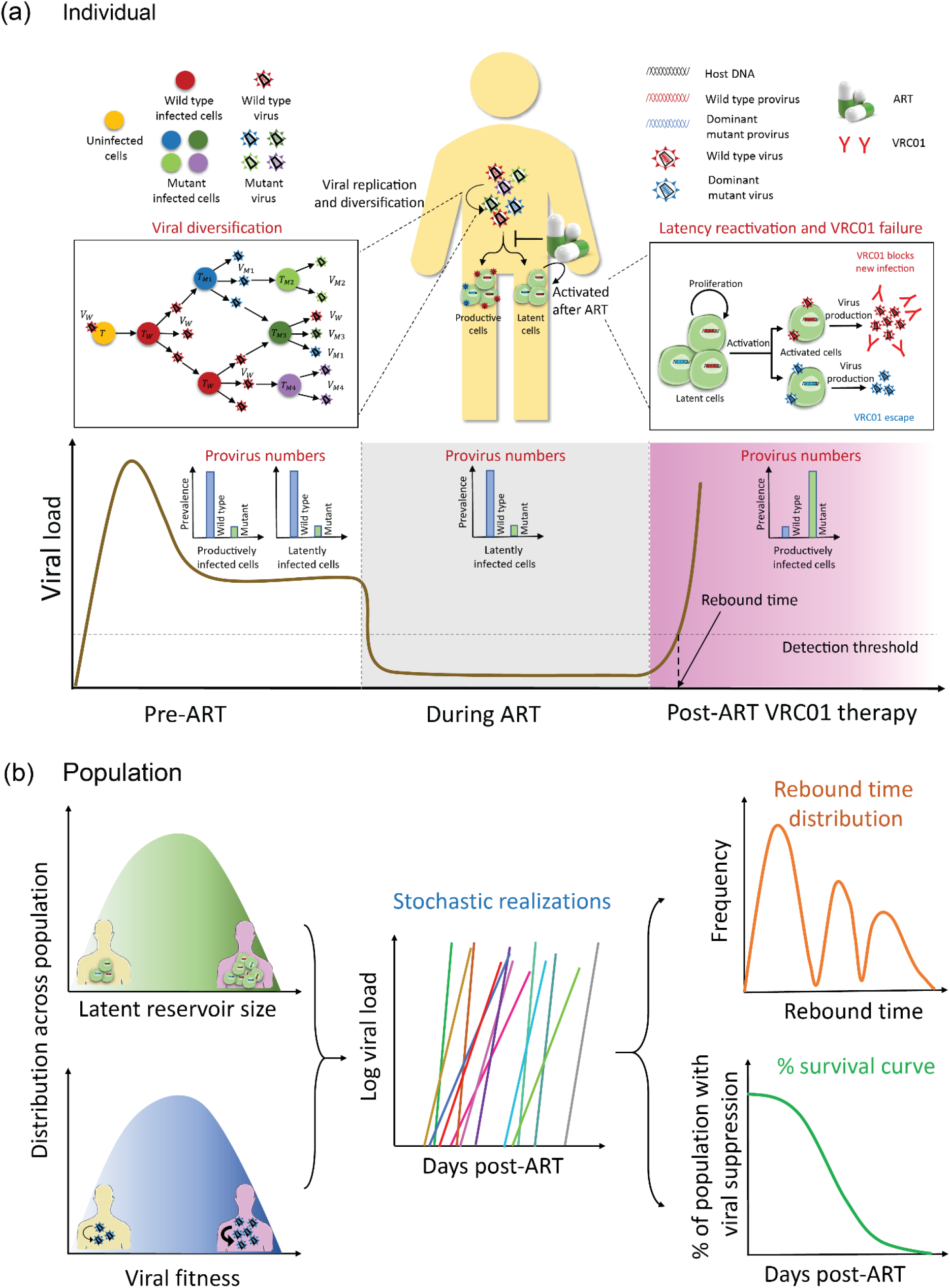
Overview of the model. **(a)** Dynamics at the individual patient level. We used a model of within-host HIV-1 evolution to estimate the pre-ART frequencies of wild-type and VRC01-resistant mutants (left), letting them be identical in productively and latently infected cells. ART eliminates the former but not the latter cells (middle). We then used a stochastic model of latent cell reactivation and viral growth to estimate the time of virological failure following VRC01 therapy (right) for a given size of the latent reservoir and the fitness of the VRC01 resistant strain. **(b)** Dynamics at the patient population level and the outcomes of clinical trials. We created a virtual patient population by sampling the latent cell pool size and mutant viral fitness during VRC01 therapy from defined distributions (left). For each individual, we performed stochastic simulations as in (a) and estimated the time to virological failure (middle), from which we obtained the distribution of breakthrough times and Kaplan-Meier survival plots (right), which we compared with clinical data.

Next, we constructed a large virtual patient population based on inter-patient variations in the size of the latent reservoir and the fitness of mutant viral strains, to reflect variations in host and viral factors, respectively, that could influence the outcomes of therapy (Fig. 1(b)). We applied our model above to each virtual patient and estimated the time of therapy failure. From these simulations, we estimated the distribution of remission times in the population and constructed Kaplan-Meier survival plots. We used data from one clinical trial to estimate the parameters defining the inter-patient variations in the virtual patient population and used them without adjustable parameters to predict the outcomes of another clinical trial, validating our model and the parameter estimates.

### Frequencies of VRC01-resistant mutants pre-existing in the latent reservoir

To estimate the frequencies of VRC01 resistant proviruses that may exist in the latent reservoir and cause VRC01 failure, we considered viral evolution before the initiation of ART in a chronically infected individual. During ART, viral replication is quickly halted^5,6^, leaving little scope for further viral diversification. We considered four mutations in the HIV-1 envelope region reported to be highly resistant to VRC01: N279K, N280D, R456W and G458D^27^. Resistance could come from strains carrying these mutations singly, in pairs, in triplets, or all together. We considered all these strains in addition to the wild-type, or VRC01 sensitive, strain in our model. The frequencies of the strains would depend on their relative fitness, which, following previous studies^54–56^, is defined in our model by two components: infectivity and replicative ability. The relative infectivity of each of the strains involved has been estimated in independent experiments^27^, which we employed (Table S1). The replicative abilities can be estimated using competitive growth assays^57^, which have been reported for some of the strains^27^. From the assays, the two fittest single mutant strains, N279K and N280D, appeared to have replicative fitness not significantly different from the wild-type^27^. We analysed data available from the assays for the other two single mutants, R456W and G458D, and the quadruple mutant^27^ using a previously developed formalism57 and estimated their replicative fitness (Fig. S1). For all the double and triple mutants, as an approximation, we set the replicative fitness to values predicted assuming zero epistasis. (This assumption is not critical to our findings, which, as we show below, depend primarily on the single mutant frequencies.) The fitness values are in Table S1. With these fitness values and all other parameters representative of chronic HIV-1 infection (Table 1), we estimated the frequencies of productively infected cells harbouring the different mutant proviral genomes (Fig. 2).

**Table 1:**
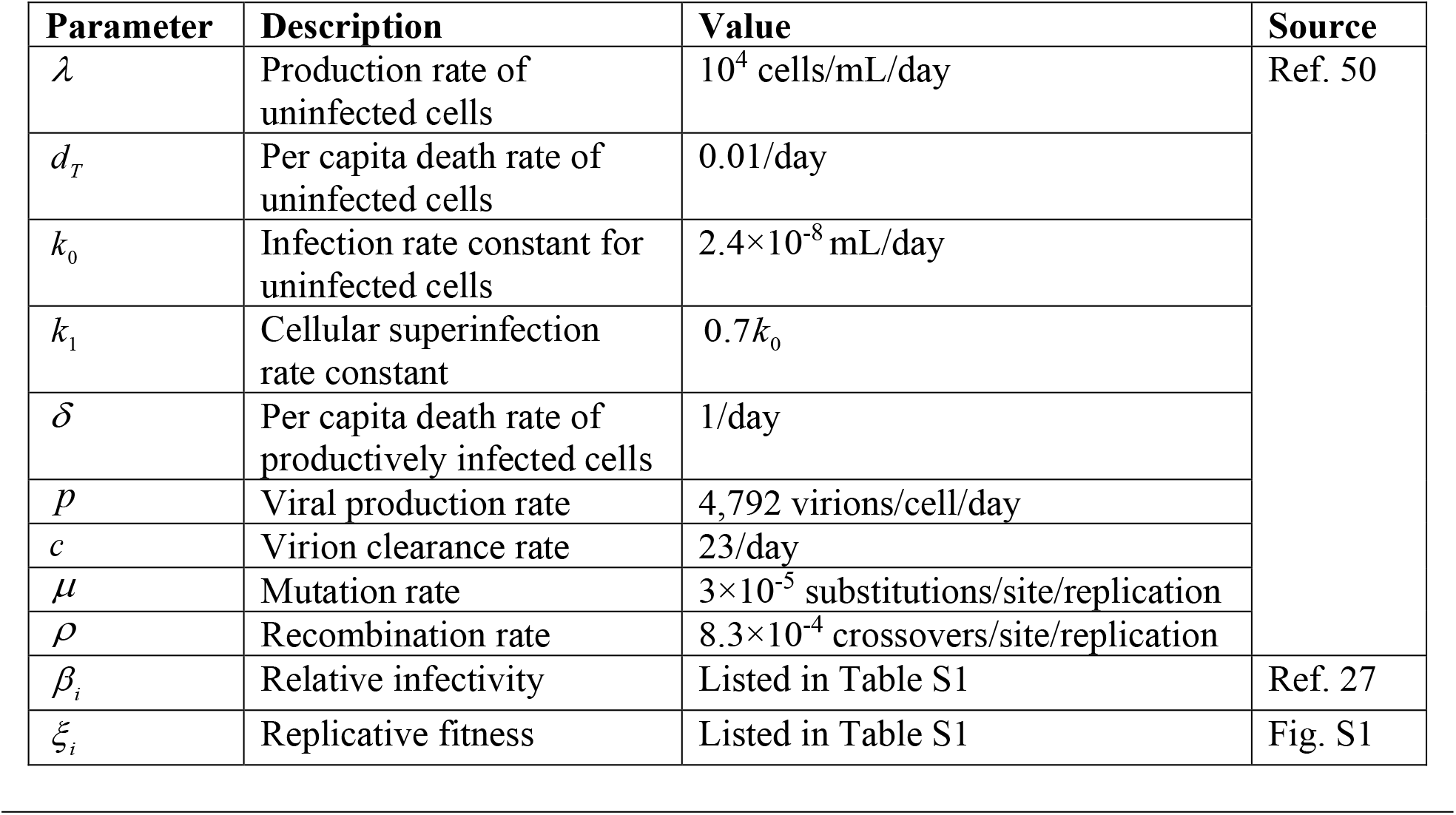
Parameters values used in the model (Eqs. 1–10) for estimating mutant frequencies.

**Figure 2:**
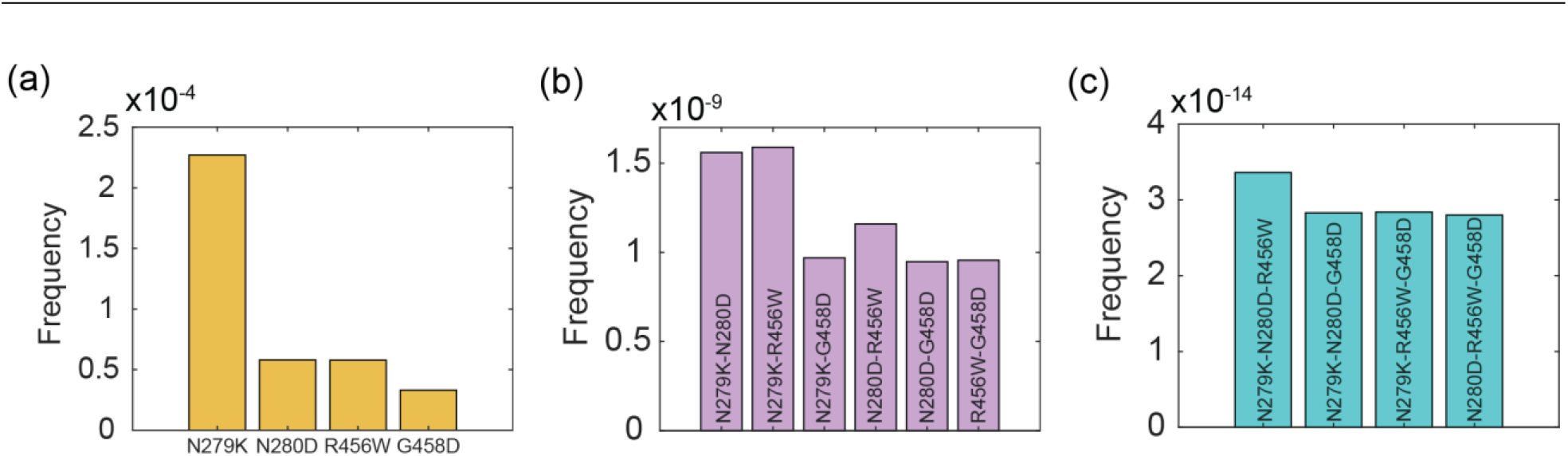
Pre-existing frequencies of mutants. Frequencies of **(a)** single mutants, **(b)** double mutants, and **(c)** triple mutants, resistant to VRC01, estimated by our model (Methods). The frequencies, including of the wild-type and the quadruple mutant, not shown here, are listed in Table S1.

Our model predicted that the fittest mutant strain, N279K, would exist at a frequency of approximately 2.3×10^−4^, the highest among the different mutants (Fig. 2(a)). Other single mutants were at frequencies ~5-fold lower. The double mutants had frequencies of ~10^−9^ (Fig. 2(b)), triple mutants of ~10^−14^ (Fig. 2(c)), and the quadruple mutant of ~10^−19^ (Table S1). These frequencies are similar but not identical to those expected from the mutation-selection balance, where a strain with *n* mutations would exist at a frequency of ~(*μ*/*a*)^*n*^, with *a* the equal per mutation fitness penalty and *μ* the mutation rate^47^. For N279K, for instance, where *a*~0.14, considering both relative replicative fitness and relative infectivity (Table S1), and with *μ*~3×10^−5^ (Ref. ^50^), the mutation-selection balance would yield a frequency of 2.14×10^−4^, slightly lower than that predicted by our model.

Following previous models, where a constant fraction of infection events is assumed to lead to latency^37,39,42^, the frequencies of the mutants are expected to be similar in productively and latently infected cells. The latent reservoir is not affected by ART directly, and even in the absence of active viral replication, the latent reservoir would decrease in size extremely slowly, taking years^7^. Thus, one could safely assume that the latent reservoir at the initiation of ART would exist nearly intact post-ART and at the start of VRC01 therapy. Given the latent reservoir size, *L*_0_, of ~10^5^−10^8^ cells in chronically infected individuals^34,39^, the expected number of cells infected with the N279K mutant proviruses would be ~23-23000. The corresponding numbers would be ~10-5000 for the other single mutants (Fig. 2). For the double and triple mutants, however, the numbers would be <0.1 and <10^−6^, respectively. On average, thus, our calculations predicted that most latently infected cells would carry wild-type, or VRC01-sensitive proviruses. A small number,
~10-10^4^ cells, would carry single mutants resistant to VRC01. Cells carrying higher mutants were unlikely to exist. The single mutants, too, may not be detectable in most cases, given current assay limits (see Introduction).

With this description of the frequencies of VRC01 resistant proviruses in the latent cell reservoir, we examined the dynamics of VRC01 failure due to latent cell reactivation.

### *de novo* generation of mutants during VRC01 therapy

From the above estimates, the lower end of the spectrum of latent cell numbers carrying single mutant proviruses, ~10 cells, is small enough that it is possible that in some individuals, due to stochastic variations in the mutant frequencies and/or infected cell numbers, the latent reservoir contains no resistant proviruses. In such a scenario, *de novo* mutation after the reactivation of latent cells carrying VRC01 sensitive proviruses would have to give rise to resistance during VRC01 therapy. Even in the presence of latent cells carrying single mutant proviruses, the large majority of cells carrying VRC01 sensitive strains may result in *de novo* mutations following reactivation of latent cells carrying VRC01 sensitive proviruses being the predominant mechanism of VRC01 failure. We developed a model to test this possibility (Text S1) and found that the estimated virological breakthrough times were far larger (over 100 days) than those observed clinically (~20 days) (Fig. S2). Although reactivation of such cells was more frequent, given their larger numbers, than the cells carrying mutant proviruses, such reactivation did not lead to lasting infection in our predictions because VRC01 successfully neutralized the viruses produced. Mutations, being intrinsically rare, did not lead to rapid enough *de novo* development of resistance. The VRC01 failure seen clinically was thus unlikely to be due to the reactivation of latently infected cells carrying VRC01 sensitive proviruses. The more likely mechanism therefore was the reactivation of latently infected cells carrying single mutant proviruses resistant to VRC01. We estimated remission times based on the latter mechanism next.

### Growth of pre-existing mutants during VRC01 therapy

We focussed on latently infected cells carrying proviruses with the N279K mutation, which, with their 5-fold higher prevalence than other mutants, were the most likely to be reactivated. Stochastic simulations (Methods) with a constant VRC01 efficacy against the mutant, indicated that the latent cell pool did not vary significantly over the durations considered (Fig. 3(a)), consistent with experiments^58^. Reactivation leading to the growth of productively infected cells carrying proviruses with the N279K mutation occurred over a duration of a few days to weeks (Fig. 3(b)). Detectable viremia, however, took longer given that the viral levels had to rise to 20 copies/ml (Fig. 3(c)). Defining the time for viremia to become detectable as the time of the failure of VRC01 therapy, or the breakthrough time, the simulations yielded a distribution of breakthrough times ranging from 20-100 d when *L*_0_ was 10^6^ cells (Fig. 3(d)), consistent with observations^17^, suggesting that virological breakthrough was likely to be due to the reactivation of cells infected latently with single mutant VRC01-resistant strains.

**Figure 3:**
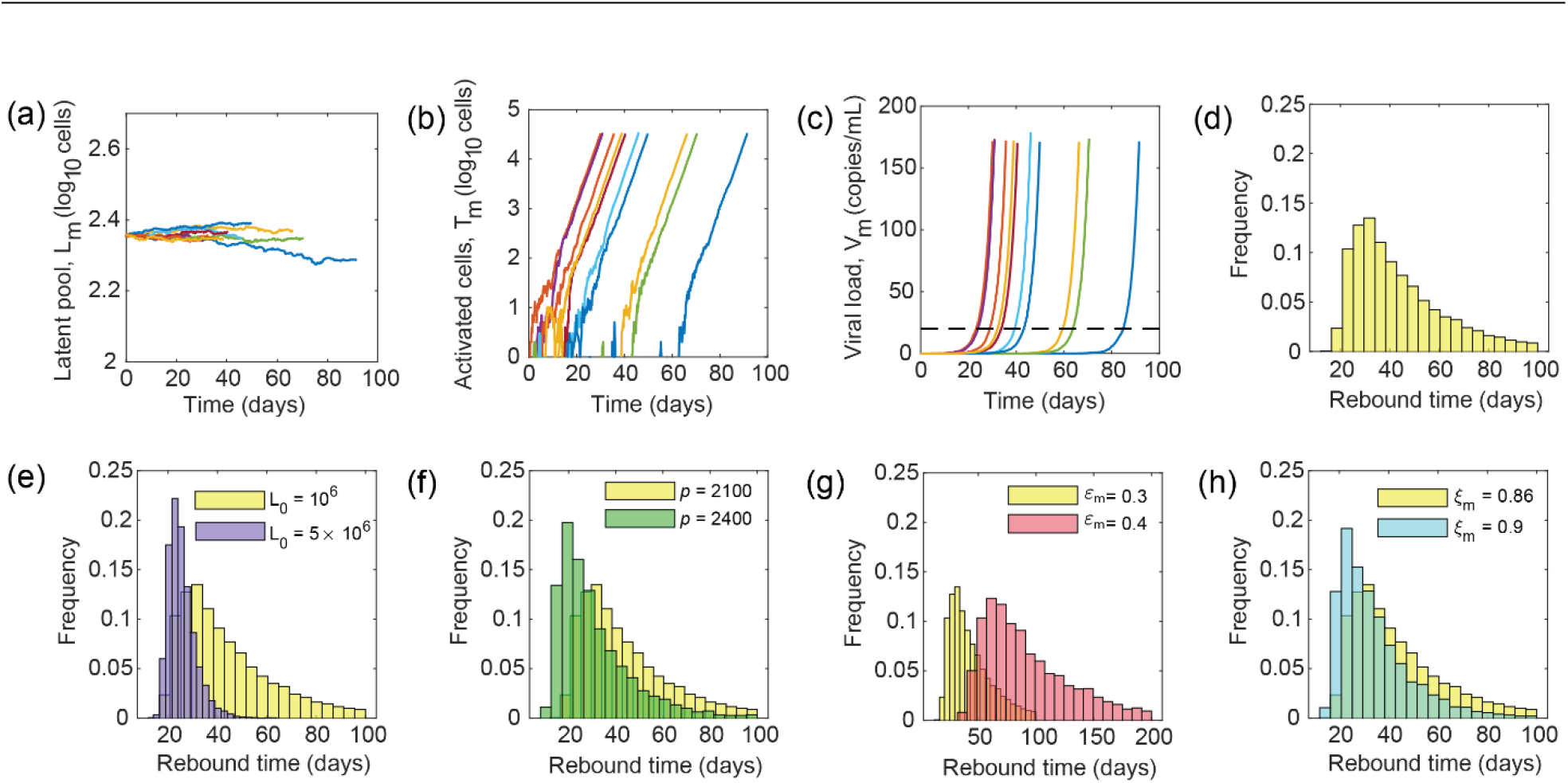
Dynamics of VRC01 failure due to pre-existing resistance. Representative trajectories of **(a)** the latent cell pool harboring resistant proviruses, **(b)** activated cells, and **(c)** VRC01-resistant viral load, obtained by our stochastic simulations (Methods). The different colors represent individual trajectories. Black dashed line shows the detection limit, crossing which marks clinical rebound. **(d)** The distribution of rebound times obtained from 5000 realizations. Here, the initial population of latently infected cells carrying the resistant mutants was set to 2.3×10^−4^×*L*_0_, where *L*0=10^6^ cells, and the VRC01 efficacy against the mutant to *ε*_*m*_=0.3. Other parameters used are in Tables 1 and 2. Variation of the distribution is shown with **(e)** initial latent pool, *L*_0_ (cells), **(f)** viral production rate (virions/cell/day), **(g)** VRC01 efficacy, and **(h)** mutant fitness.

The breakthrough time depended on the time for latent cell reactivation as well as for the subsequent establishment of successful infection. Thus, higher *L*_0_, which decreased reactivation times, led to earlier breakthrough (Fig. 3(e)). Increasing the viral production rate (Fig. 3(f)), lowering drug efficacy ((Fig. 3(g)) or increasing viral fitness (Fig. 3(h)), which improved the chances of establishment of successful infection after reactivation, all led to more rapid virological breakthrough. These factors are likely to vary across individuals. We examined next how their influence would manifest in clinical trials and whether our simulations could capture the VRC01 failure seen in the trials.

### Multimodal distribution of breakthrough times in a virtual patient population

We focused here on the A5340 trial^17^ where VRC01 therapy was initiated a week before the end of ART on patients with well-controlled viremia. 3 doses of VRC01 were administered, with a gap of 3 weeks between successive doses. Virological breakthrough was detected when the viral load crossed 20 copies/ml. To mimic the trials, it was necessary not only to account for inter-patient variations but also to consider time-varying efficacies of drugs within an individual, which can significantly affect the development of drug resistance^49,59^. We therefore first considered VRC01 pharmacokinetics and fit a model of biphasic decay to data of the VRC01 plasma concentration profile following a single intravenous dose^25^ (Fig. 4(a) inset). Using the resulting best-fit parameters, we predicted the multiple dose pharmacokinetics for the dosing protocol above and estimated the time-varying efficacy, *ε*_*m*_ (Fig. 4(a)), which we employed in our stochastic simulations.

**Figure 4:**
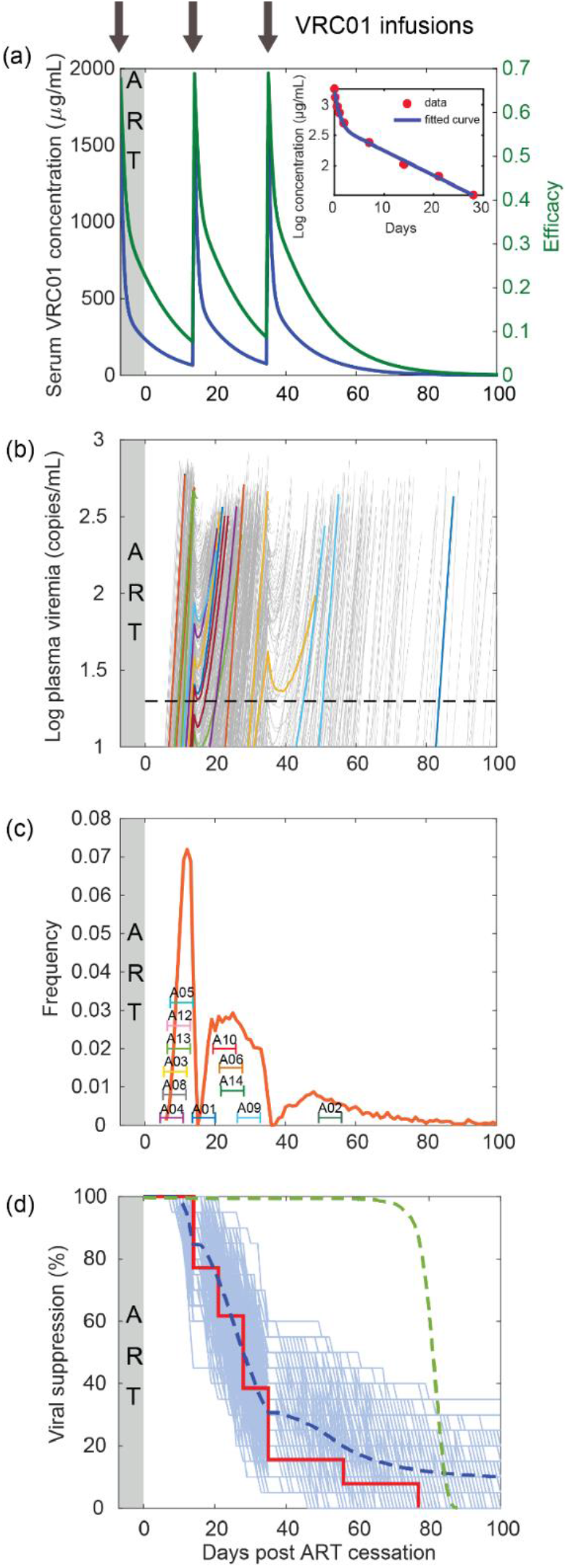
Recapitulating VRC01 failure in the A5340 trial. **(a)** Fits of our model of single-dose plasma VRC01 pharmacokinetics (line) to data^25^ (symbols) (*Inset*). Best-fit parameter estimates are in Table 2. Corresponding multiple dose concentration profiles (blue) and the VRC01 efficacy against a mutant strain with *IC*_50_=800 *μ*g/mL (green) are shown. Black arrows indicate VRC01 infusions. **(b)** Stochastic realizations of the dynamics in a virtual population of 20000 patients, manifesting as changes in plasma viremia. Each grey line represents a virtual patient. Some randomly chosen trajectories are colored to aid visualization of the dynamics. Note that ART was continued after the first infusion of VRC01 for 1 week. The detection limit of 20 copies /mL is marked as a black dashed line, crossing which marks clinical viral rebound. **(c)** The corresponding distribution of rebound times (orange). Rebound times of the participants in the A5340 trial with a 1-week uncertainty period, representing the gap between successive viral load measurements, are also marked. **(d)** Kaplan-Meier plot for the A5340 trial based on the percentage of patients with viremia >200 copies/mL. Each light blue line is a survival curve generated by randomly choosing 20 patients from the virtual population above. The dashed blue line is the mean of all these survival curves. The data from the trial is shown as a red solid line. The green dashed line shows the survival plot when failure occurs because of wild type or sensitive strains.

Next, we created a virtual patient population to mimic inter-patient variations expected in clinical trials. A number of host factors, including HLA types, are known to affect the ability of individuals to control viremia^60^. Similarly, viral factors have also been argued to determine the level of viremia in chronic infection^61^. A convolution of host and viral factors is expected to determine clinical outcomes. The specific factors involved and how they vary across individuals, however, is not fully established^46,60^. Here, we employed a parsimonious approach where we let two parameters, one reflective of variations in host factors and the other viral factors, define the inter-patient variations. We thus constructed a virtual patient population with different initial latent pool sizes, *L*_0_, subsuming variations in all host-factors, and viral production rates, *p*, subsuming variations in all viral factors. For each individual, we sampled *L*_0_ and *p* independently from pre-defined distributions (Methods) and ran a stochastic simulation to describe the ensuing dynamics resulting in virological breakthrough.

The simulated dynamics showed breakthrough times varying from a few days to a few months post cessation of ART across individuals (Fig. 4(b)). Following breakthrough, the viral load rose sharply. It was suppressed partially following the administration of a VRC01 dose, but then rose again once the VRC01 level fell. The patterns were similar to the viral load resurgence patterns seen in patients^17^. From the breakthrough data, we estimated the distribution of virological breakthrough times in this virtual patient population (Fig. 4(c)). We found that the distribution was multimodal. Until a week or so following the cessation of ART, no breakthrough was expected based on the distribution because of high VRC01 levels in circulation. As the VRC01 concentration declined, breakthrough began. The distribution of breakthrough times peaked when the VRC01 concentration was at its trough level, just before the administration of the second VRC01 dose. Following dosing, the distribution dropped steeply, and rose again as the VRC01 concentration waned. It then attained a peak that was smaller than the first peak and began to fall subsequently. The smaller size of the peak and the subsequent fall was due to a convolution of the effect of VRC01 and the natural distribution of breakthrough times in the absence of treatment. Post ART cessation, it has been shown that the distribution of mean rebound times is unimodal and declines following its peak^41^. In other words, after the peak, the population of individuals that suffers breakthrough decreases with the breakthrough time. In our simulations, this decline is what explains the smaller second peak compared to the first and the drop in the distribution after the second peak although the VRC01 levels were low. Of course, with the third dose, the distribution dropped sharply again due to the rise in the VRC01 level and then rose as the VRC01 level waned. The distribution attained a third peak that was even smaller than the second peak and ended in a long tail representing the small fraction of individuals who experienced longer remission times than studied in our simulations.

Based on the distribution, we constructed a Kaplan-Meier survival plot, which at each time point marked the percentage of the virtual patient population that was still under remission, defined as viremia <200 copies/mL^17^. The plot, as expected, indicated no failure for a short period, ~1-2 weeks, after ART, then dropped sharply, reaching 50% failures in about 4 weeks, and displayed a long tail with a small percentage, ~10%, maintaining remission for longer than the duration of our simulations (100 days) (Fig. 4(d)). We examined next whether these predictions could recapitulate clinical observations.

### Model calibration to recapitulate the A5340 trial

Comparing our simulations with data required knowledge of the distributions of *L*_0_ and *p* in patients. We chose *p* to mimic viral growth rates after rebound. The growth rate can vary from 0.4-1.5/day^39,42^. We therefore chose the mean value of *p* to yield a growth rate of 1/day. Further, we let it follow a normal distribution with a standard deviation that ensured positive growth rate across two standard deviations from the mean. The resulting distribution of *p* is mentioned in Table 2. Although estimates of the variations in *L*_0_ exist^34,39^, in our simulations, they must accurately mimic the variations in the pool carrying resistant proviruses, which are not known. We therefore adopted the following approach. We decided to employ data from the A5340 trial to estimate parameters characterizing the distributions of *L*_0_ and then validate them using an independent trial, the NIH trial. Both the trials involved small sample sizes, ~10 patients^17^. Non-linear mixed effects modeling, designed particularly to estimate parameter distributions using clinical data from small sample sizes, works with deterministic but not stochastic models^62,63^. Consequently, we adopted a heuristic approach to estimate the distributions of *L*_0_. We recognized that the product *a_c_L*_0_, with *a_c_* the latency reactivation rate, determines the waiting time for viral recrudescence; the larger the product, the earlier would be the reactivation of the latent reservoir. We fixed *a_c_* based on previous estimates^37,39^. (Note that previous studies report a wide range for *a_c_*.^35,39,42^) Through small test simulations, we identified approximate values of *L*_0_ that mimicked the mean waiting times seen in patients in the A5340 trial. We then performed more detailed simulations by varying the distribution of *L*_0_ around the approximate parameters and identified those distributions that best described the Kaplan-Meier survival data from the A5340 trial (Fig. S3). The resulting parameters are also in Table 2. Similarly, we explored the implications of variations in the *IC*_50_ of VRC01 against the mutant and found that values ≥800 μg/mL captured the data well (Fig. S4).

**Table 2:** Parameters values used in the model of viral rebound (Eqs. 11–20). Parameters not listed are in Table 1.

| Parameter | Description | Value | Source |
| --- | --- | --- | --- |
| $L_0$ | Initial latent reservoir size | $Log_{10}(L_0) \sim N(6.3, 0.43)$ cells (lognormal distribution) | Fig. S3 |
| $\rho_l$ | Per capita proliferation rate of latently infected cells* | 0.0042/day | Refs. 37,39 |
| $d_l$ | Per capita death rate of latently infected cells* | 0.004/day | Ref. 37 |
| $a_c$ | Per capita activation rate of latently infected cells* | $7.8 \times 10^{-4}$ /day | Refs. 37,39 |
| $f$ | Fraction of new infections that lead to latency | $10^{-6}$ | Ref. 37 |
| $k$ | Infectivity<sup>\$</sup> | $1.6 \times 10^{-12}$ /day | Ref. 50 |
| $p$ | Viral production rate | $p \sim N(2100, 200)$ virions/cell/day (normal distribution) | Refs. 39,42 |
| $U$ | Uninfected cell population<sup>\$</sup> | $1.5 \times 10^{10}$ cells | Ref. 50 |
| $\varepsilon_{ART}$ | ART efficacy | 0.99 | Assumed |
| $IC_{50,m}$ | 50% inhibitory concentration for the N279K mutant<sup>&</sup> | 800 μg/mL | Fig. S4 |
| $A_1$ | Portion of VRC01 dose associated with first phase decay<sup>#</sup> | 451 (320, 580) μg/mL | Best-fit (Fig. 4(a) inset) |
| $A_2$ | Portion of VRC01 dose associated with second phase decay<sup>#</sup> | 1253 (910, 1596) μg/mL | |
| $\eta_1$ | First phase decay rate<sup>#</sup> | 0.093 (0.078, 0.109) /day | |
| $\eta_2$ | Second phase decay rate<sup>#</sup> | 1.374 (0.752, 1.995) /day | |

\* $\rho_l$ , $a_c$ , and $d_l$ together yield a half-life of ~40 months for the latent pool, consistent with experiments<sup>7</sup>.
<sup>\$</sup>Corresponding to 15 L of body fluid volume.
<sup>&</sup>Estimates indicate a value much larger than 50 μg/mL<sup>27</sup>, but a precise value is lacking.
<sup>#</sup>95% confidence limits are in brackets.

Simulations with the resulting distributions recapitulated data from the A5340 trial in two ways. First, we found that 11 of the 12 patients who experienced treatment failure despite high concentrations of VRC01 in circulation^17^ had breakthrough times close to the peaks in the multimodal distribution of breakthrough times we predicted (Fig. 4(c)). The number of patients with breakthrough times associated with the different peaks was also proportional to the area under the peaks. Note that the area under a peak is a measure of the probability, and hence the frequency, of failure corresponding to the times associated with the peak. 6 patients failed during the times associated with the first peak, 4 with the second peak, and one with the third peak. The areas under the peaks from our simulations yielded failure percentages of 35%, 44%, and 21%, respectively. The first two peaks, thus, appeared to have comparable failure percentages, whereas the third was substantially smaller. Given the small sample size in the clinical trial, the distribution of patients into the three peaks appeared to be consistent with the estimated failure percentages. One patient (A01) appeared to fail at the trough in the distribution after the first peak, and this could be due to stochastic effects or variations not captured in our virtual population.

Second, Kaplan-Meier plots based on breakthrough times, *i.e.*, times for the viral load to reach 200 copies/ml, from the virtual patient population were in close agreement with the clinical data (Fig. 4(d)). Here, to account for the small sample size in the trials, we chose many samples of 20 individuals each, selected randomly from our virtual patient population, and constructed Kaplan-Meier curves for each sample. The clinical data fell within the ranges defined by these curves. Further, the mean of these curves was in close agreement with the data. Accordingly, 50% of the treated population exhibited a breakthrough time of >3 weeks from the end of ART (or 4 weeks from the start of VRC01 therapy), consistent with the clinical data. The simulations tended to over-predict the clinical data for long remission times (>40 d). We attributed this to the presence of individuals with strong immune responses and/or small latent cell populations, including post-treatment controllers^60^, who may not be seen in the small population of 12 individuals in the A5340 trial. Secondly, after the effect of VRC01 wanes, failure may also occur due to the reactivation of wild-type virus. Indeed, simulations with the wild-type virus alone (Text S1) showed that once the VRC01 level declined sufficiently, breakthrough with the reactivation of wild-type virus would become more likely than with the mutant virus (green dashed line in Fig. 4(d)). Our simulations, thus, successfully recapitulated the data from the A5340 trial.

### Model validation with the NIH trial

To test and validate our model and the parameter estimates, we applied our simulations to describe a second, independent clinical trial, the NIH trial^17^, where 10 individuals were subjected to VRC01 therapy during an ATI. Treatment commenced 3 days before ART cessation. Subsequent doses were administered on weeks 2 and 4 and then every month until 6 months. To describe the resulting breakthrough data, we created a virtual population exactly as above and subjected it to therapy following the clinical protocol. VRC01 pharmacokinetics was predicted based on the corresponding dosing times (Fig. 5(a)). All the parameters were kept the same as those as in our simulations of the A5340 trial.

**Figure 5:**
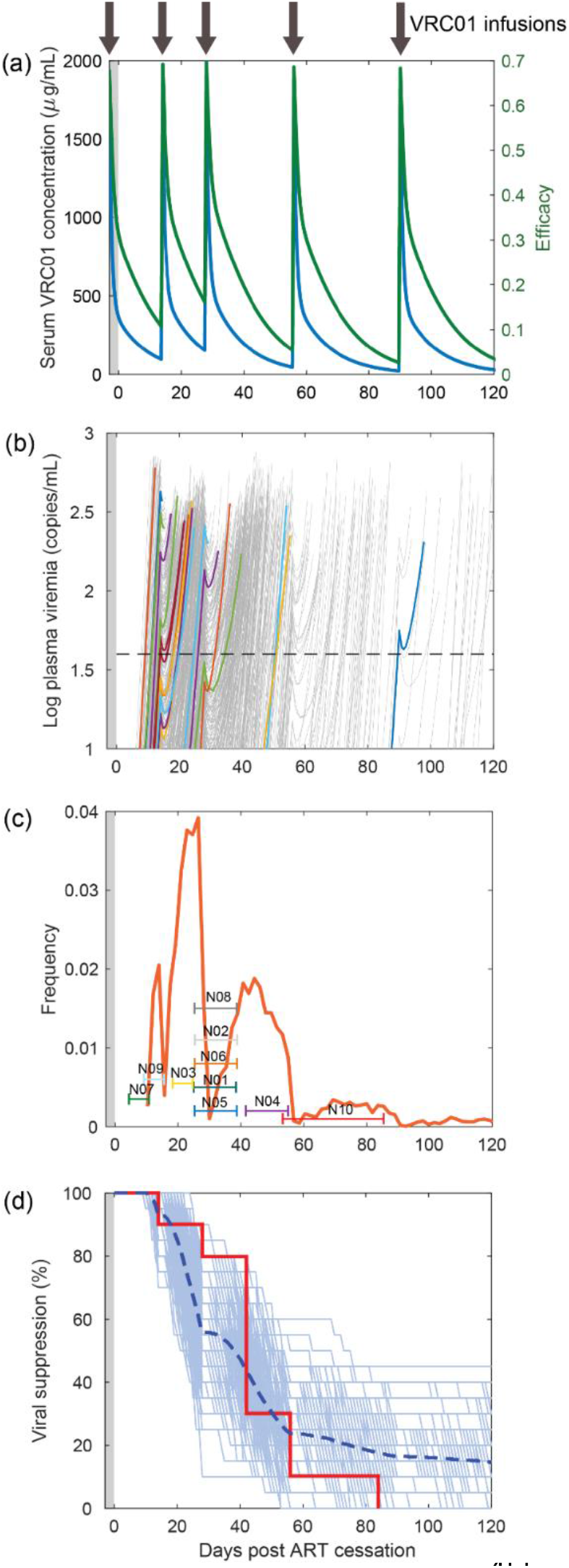
Recapitulating VRC01 failure in the NIH trial. **(a)** Multiple dose concentration profiles (blue) and the VRC01 efficacy against a mutant strain with *IC*_50_=800 μg/mL (green). Black arrows indicate VRC01 infusions. **(b)** Stochastic realizations of the dynamics in a virtual population of 10000 patients, manifesting as changes in plasma viremia. Each grey line represents a virtual patient. Some randomly chosen trajectories are colored to aid visualization of the dynamics. ART was continued after the first infusion of VRC01 for 3 days (grey shaded region). The detection limit of 40 copies /mL is marked as a black dashed line, crossing which marks clinical viral rebound. **(c)** The corresponding distribution of rebound times (orange). Rebound times of the participants in the NIH trial are marked along with their uncertainties based on measurement frequencies. **(d)** Kaplan-Meier plot for the NIH trial. Each light blue line is a survival curve generated by randomly choosing 20 patients from the virtual population above. The dashed blue line is the mean of all these survival curves. The data from the trial is shown as a red solid line.

We found that viral rebound trajectories were similar to those in the A5340 trial, with sharp rises in viral load following breakthrough and wide inter-patient variations (Fig. 5(b)). The distribution of breakthrough times again followed a multimodal distribution with the peak widths broadening progressively with each dose and culminating in a long tail (Fig. 5(c)). Based on the areas under the peaks, the percentages of failure at times corresponding to the four peaks were 7%, 37%, 32%, and 24%, respectively. Indeed, in agreement, 7 of the 10 patients failed at times corresponding to the second and third peaks. 2 patients were associated with the first peak and one with the last. More precise timing of failure of the patients was not possible given the much larger intervals between successive viral load measurements in the NIH trial compared to the A5340 trial. Nonetheless, the distribution of failure times between peaks, given the small sample size, was remarkable. Furthermore, Kaplan-Meier plots captured the clinical data quantitatively (Fig. 5(d)), indicating that our simulations accurately mimicked the response of patients to VRC01 therapy in the NIH trial.

The difference between the A5340 trial and the NIH trial was in the dosing protocol alone. Our simulations captured both these datasets by simply changing the dosing protocol accordingly, and required no other adjustments, thereby providing a strong test and validation of our model. The model may be applied to predict the outcomes of other possible dosing protocols, which could involve changing dosages or half-lives (Fig. S5), providing a framework for rational therapy optimization.

## DISCUSSION

Passive immunization with HIV-1 bNAbs holds promise as a strategy to achieve long-term remission of HIV-1 infection^20,64^. Following its success in macaques^20,21^, enormous efforts are underway to translate it to humans^65^. Trials with VRC01 administered to chronically infected patients following the cessation of successful ART saw rapid virological failure despite the presence of suppressive concentrations of VRC01 in circulation^17^, signaling a potential vulnerability of such bNAb therapies. Here, using mathematical modeling and analysis of clinical data, we elucidated the likely cause of this rapid VRC01 failure. Model predictions attribute this failure to the reactivation during therapy of cells latently infected with VRC01 resistant proviruses before ART initiation. To arrive at this inference, we constructed a mathematical model that integrated within-host viral evolution, latency reactivation, and viral dynamics with VRC01 pharmacokinetics, and applied it to simulate the outcomes of therapy in a virtual patient population. Our simulations recapitulated data from two independent clinical trials^17^, giving us confidence in the inference. Accounting for pre-existing resistant strains in the latent reservoir would be important to the success of bNAb-based therapies.

That mutation-driven resistance can be an important cause of bNAb failure has been recognized earlier^27–30^. Indeed, efforts are ongoing to identify bNAbs, including in the VRC01 class, that are less vulnerable to such resistance.^66^ For bNAb therapies that target maintenance of HIV remission post ART cessation during chronic infection, our study prescribes the requisite genetic barrier to resistance, *i.e.*, the number of mutations HIV-1 must accumulate to develop significant resistance to the therapy. Using a detailed description of within-host HIV-1 evolution, our study predicts that with the estimated latently infected reservoir size of 10^5^-10^8^ cells in a chronically infected individual^34,39^, proviral strains carrying single but not more resistance mutations would pre-exist in the latent reservoir. Consequently, a genetic barrier of 2 or more would ensure that pre-existing resistance would not compromise therapy. A bNAb with a genetic barrier of one, like VRC01, would be predestined to fail, as was observed in clinical trials^17^. When used in combination with another bNAb, however, the overall barrier would cross the threshold of 2, diminishing the chances of such failure. Resistance would then have to develop by *de novo* mutation during therapy, which according to our model would be unlikely as long as adequate bNAb concentrations exist in circulation and restrict the replication of the wild-type virus. Further, by explicitly incorporating bNAb pharmacokinetics, our model provides a framework with which optimal dosing protocols could be identified that would ensure adequate bNAb concentrations throughout.

Early ART initiation has been argued to restrict the viral reservoir size and improve the chances of post-treatment control^10,37^. Our study predicts an additional advantage of early ART initiation: The chances of failure due to resistance are reduced. This reduction happens in multiple ways. First, most HIV-1 infections involve a single founder virus, which gradually evolves into the diverse quasispecies seen in chronic infection.^52,67,68^ Thus, the frequency of mutants at the time of ART initiation is expected to be smaller, the earlier the initiation.^52^ Second, given the long half-life of the latent reservoir^7^, cells infected early during infection are likely to exist in the latent reservoir in a much higher proportion than in the actively replicating or plasma compartments. Given the limited viral diversity early in infection, the latent reservoir is likely to have a much higher representation of bNAb-sensitive strains than estimated using our model of chronic infection. Finally, if the reservoir size is restricted, due to early ART initiation, far fewer mutant proviruses, or none at all, may exist in the latent reservoir. Together, thus, early initiation of ART is expected to reduce the chances of bNAb failure. Indeed, in trials where ART was initiated early, in the acute phase of infection, no genotypic resistance to VRC01 was detected from sequence analysis of viral strains post failure^69^. Similarly, a combination of two bNAbs, including one in the VRC01 class, administered early in infection saw no resistance to therapy in SHIV-infected macaques.^20^ Surprisingly, however, the breakthrough times for VRC01 therapy following ART in acute infection were found to be similar to those seen in the trials in chronic infection we studied.^17,69^ Additional mechanisms that are not predominant in the chronic phase thus appear to cause VRC01 therapy failure in individuals treated with ART early. One possibility could be that CD8 T cell exhaustion is weaker and/or more reversible in the acute phase than in the chronic phase because of the much longer duration of antigen exposure in the latter scenario.^70,71^ The greater associated immune activation levels in the acute phase could imply more rapid reactivation of latently infected cells, which could offset the advantage from the lower frequencies of mutants. To test this possibility, models that incorporate CD8 T cell exhaustion^37,72–75^ would have to be integrated with our model of stochastic latency reactivation, a promising avenue for future study.

An important question in HIV cure research is how early should ART be initiated to maximize post-treatment control^60^. While starting early would restrict the reservoir size and improve the chances of post-treatment control, starting it too early would not allow enough time for the development of an immune response, compromising post-treatment control. If bNAb therapy were to be used post-ART to improve the chances of post-treatment control, the development of resistance to bNAbs would have to be factored in along with the latter trade-off between reservoir size and immune response strength to arrive at the optimal timing of ART initiation. Our study provides a framework that could be used to test whether pre-existing resistance in the latent reservoir would compromise therapies with other bNAbs, especially those belonging to the VRC01-class^76–78^, which are under trial for both preventive^24,79,80^ and therapeutic^20^ vaccination and are known to fail via diverse mutation-driven resistance pathways^28^.

An early modeling study compared the likelihood of the failure of therapy due to pre-existing resistance versus *de novo* generation of mutants in the context of ART and found that ART was more likely to fail due to pre-existing resistance.^48^ A more recent modeling study examined the likelihood of the failure of different antiretroviral drug combinations and explained how patient adherence influences such failure.^53^ Our findings are consistent with these studies and show that bNAbs too are vulnerable to pre-existing resistance, but the resistance now is restricted to the latent reservoir. Because the reservoir is small compared to the pool of infected cells pre-ART, the required genetic barrier for bNAb therapies is estimated to be lower than for ART. Thus, a combination of 2 bNAbs is predicted to overcome resistance, whereas first-line ART necessarily contains 3 drugs.

Previous studies have used either fully deterministic or fully stochastic frameworks to describe treatment failure^48,49,53^, making their predictions approximate or computationally expensive. Here, we devised a strategy that retained both accuracy and computational tractability. We used a deterministic framework to estimate the frequencies of mutant proviral genomes in the latent reservoir pre-treatment and then a stochastic framework to estimate latency reactivation and treatment failure times. The deterministic framework was shown previously to agree well with stochastic population genetics-based simulations^50^, giving us confidence in the strategy. In the stochastic simulations, we considered only latently infected cells containing the dominant resistant provirus, which accurately described the development of drug resistance without requiring expensive computations. Indeed, we were able to capture two independent clinical trials of VRC01 failure with this hybrid framework, reiterating its applicability. Such hybrid deterministic-stochastic frameworks have been successfully applied in other settings, such as in predicting the pre-existing frequency of hepatitis C virus strains resistant to drugs^81^, but not, to our knowledge, with HIV-1 infection.

The sample sizes involved in the clinical trials we studied were small, ~10 patients each^17^. Nonlinear mixed-effects models have been used in recent studies to infer the effects of interventions by analyzing data from such trials^82^. For instance, bNAb therapy was argued to improve immune responses 8-fold over controls and not to synergize with the TLR7-agonist^82^. The strength of such an approach lies in its robust parameter estimation and the ability to infer effects at the population level using data from small sample sizes. The approach, however, does not work when the underlying model is stochastic, which is the case in our study. Also, when the effects are explicitly parameterized, as is often done^82,83^, their quantitative estimates are restricted to the specific conditions studied. Thus, for instance, how a change in the dosage or dosing protocol would alter the effect becomes difficult to predict. Our approach, being fully mechanistic, is not similarly limited. Indeed, with the parameters that captured the A5340 trial, by simply changing the dosing protocol, our model captured data from the NIH trial without any adjustable parameters. Our model could therefore be used potentially to comparatively evaluate alternative treatment protocols and suggest optimal ones.

We recognize that in the trials we considered^17^, the dominant mutant seen post treatment failure was not the same across individuals. The mutations, however, were all typically concentrated in the same genomic regions^17^, suggesting that structural or conformational modifications that could drive VRC01 resistance could be produced by many mutations in the same genomic region. While we have focused on the most frequent mutant as it is the most likely to have caused resistance, and as has been done in earlier studies^53^, stochastic variations could result in latent cells carrying other mutants being reactivated and reestablishing infection. The mutants selected could also differ based on inter-host variations and the genetic backgrounds of the infecting strains. Our simulations must thus be viewed as reflecting therapy failure arising from the dominant mutant which could differ across hosts and which in our study is accounted for by the inter-host variation in viral fitness.

bNAbs are known to engage the immune system via multiple mechanisms^20,84–87^. The result could be heightened activation, which for HIV-1 infection, could mean greater susceptibility of target CD4 T cells. At the same time, it could imply greater reactivation rates of latently infected cells. Indeed, the latency reactivation rates employed in our study were higher than those estimated for historical controls, the latter based on viral recrudescence post ART and in the absence of further intervention^39^. When bNAb therapy is administered post ART, cells latently infected with wild-type strains, which would be in a vast majority, would get frequently reactivated and produce virions but without causing sustained infection. The virions produced, being bNAb sensitive, would get neutralized and cleared by the bNAbs. Previous studies have argued that in the process bNAbs can stimulate CD8 T cells^20,87^, improving the overall immune activation status, possibly explaining the higher latency reactivation rate we required to describe VRC01 failure than previously used to describe historical controls. Our parameters would thus tend to underpredict the remission times in the historical controls. Previously, too, differences in the viral growth rates before and after ART have been assumed in order to capture viral recrudescence accurately and have been attributed to different strengths of the immune responses in the respective periods.^39^ Mechanisms that could explain the differences in these latency reactivation rates are yet to be identified. A unified framework that describes remission both with and without bNAb therapy awaits future studies that would quantify how bNAbs influence immune responses.

Non-nucleoside reverse transcriptase inhibitors (NNRTIs), administered as part of ART, have been found to delay breakthrough post ART^23^. Patients administered NNRTIs were excluded from the A5340 trial and were switched to an integrase-inhibitor based regimen 2 weeks before VRC01 therapy in the NIH trial to eliminate the confounding effects of NNRTIs. We therefore did not consider the effect of NNRTIs in our study. A number of host^60^ and viral^61^ factors are thought to be involved in determining disease progression and treatment outcomes. Our model subsumed inter-patient variations in these factors into variations in two factors, the latent pool size and the viral production rate. Virtual patient populations that we created based on variations in these minimal factors captured clinical data from two independent clinical trials, justifying the approximation, and suggesting that a small subset of factors may be adequate to capture outcomes of such bNAb therapies. Future studies may consider the variations in other factors explicitly to ascertain the validity of the approximation, especially for other kinds of bNAb-based interventions.

In summary, our study presents an explanation of the failure of VRC01 therapy to sustain HIV remission post ART, captures clinical data quantitatively, highlights the importance of accounting for pre-existing resistance in designing effective bNAb-based therapies, and facilitates rational optimization of such therapies.

## METHODS

We considered the scenario where a chronically infected individual maintains viral suppression with ART and is then subjected to VRC01 therapy concomitantly with cessation of ART. Because active viral replication is halted by successful ART, virological breakthrough during VRC01 therapy must arise from the reactivation of latently infected cells. We developed a model to estimate the timing of the failure of VRC01 therapy via the growth of resistant viral mutants pre-existing in the latent reservoir (Fig. 1). Our approach was to estimate the population of latently infected cells carrying the resistant strains and then to follow their reactivation leading to successful infection. We then considered a virtual patient population subjected to the same therapy and applied our model to recapitulate data from clinical trials.

### Mathematical model of virological breakthrough in a single infected individual

#### Within-host HIV-1 evolution and the frequencies of pre-existing resistant strains

To estimate the population of cells latently infected with VRC01-resistant proviral genomes, we reasoned that the frequencies would be the same as in productively infected cells before treatment initiation because the probability that a particular cell becomes latently infected is not known to depend on the nucleotides at the VRC01 resistance loci. We estimated the frequencies of VRC01-resistant strains in productively infected cells using an approach developed previously to quantify the pre-existence of resistance to antiretroviral drugs^50^. The approach combines virus dynamics with evolution, incorporating mutation, cellular superinfection, recombination, and fitness selection. We present details below.

#### Virus dynamics

We considered *n*=4 positions in the *env* region where mutations with high level resistance to VRC01 have been identified, namely N279K, N280D, R456W and G458D^27^. The time evolution of the populations of cells and virions containing the different single, double, triple, and quadruple mutants were described using the following equations:

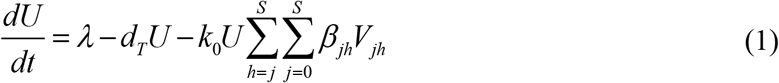

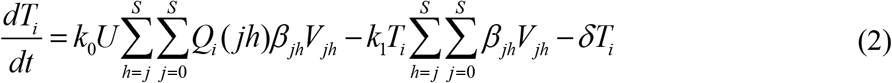

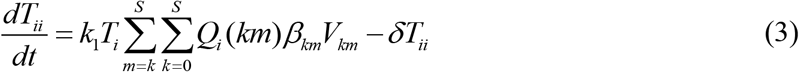

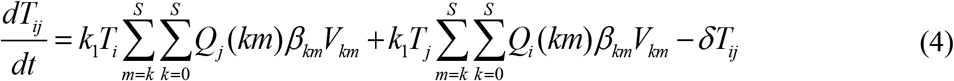

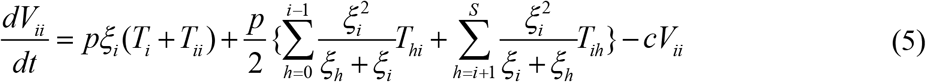

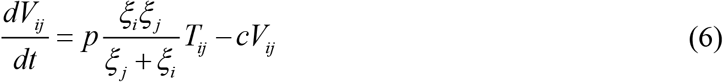

Here, uninfected target CD4+ T cells, *U*, are produced from the thymus at the rate *λ* and are lost at the per capita death rate *d*_*T*_. They are infected by virions *V*_*jh*_ containing genomes *j* and *h* with the second order rate constant *k*_0_*β*_*jh*_, where *k*_0_ is the infectivity of wild-type virions and *β*_*jh*_ is the relative infectivity of virions *V*_*jh*_. We assumed that *β*_*jh*_=(*β*_*j*_+*β*_*h*_)/2, where *β*_*j*_ is the infectivity of genome *j* relative to the wild-type. The indices *j* and *h* denoting viral genomes range from 0 to *S* = 2^*n*^ −1. Thus, *S*=15 here, with the different genomes including the wild-type (or sensitive strain), the 4 single mutants, 6 double mutants, 4 triple mutants, and the quadruple mutant, amounting to a total of 16 types. We denoted the genomes serially, starting with ‘0’ for the wild-type and ending with *S* for the quadruple mutant. Each viral particle contains two genomes, not necessarily identical. The number of distinct viral particle types, *V*_*jh*_, where *j*∊{0,1,..,*S*} and *h*∊{*j*,*j*+1,..,*S*}, is thus (*S*+1)(*S*+2)/2, which here would be 136. (The range of values of *h* is to ensure that virions *V*_12_ and *V*_21_, for instance, which are identical, are not counted separately.) Summing over all these viral types yields the total rate of loss of *U* due to infections in Eq. 1.

Following infection with a virion *V*_*jh*_, reverse transcription, which includes mutation and recombination, yields genome *i* with a probability *Q*_*i*_(*jh*), where *i*∊{0,1,..,*S*}. On average, thus, a fraction *Q*_*i*_(*jh*) of the infections with virions *V*_*jh*_ yield productively infected cells carrying single proviruses *i*, which we denote *T*_*i*_. Summing over all *V_jh_* yields the total rate at which cells *T_i_* are produced. These cells die at the per capita rate *δ*. They can also be infected again, but with a lower infectivity *k*_1_ because infected cells downregulate their CD4 receptors, rendering further infections difficult^88,89^. The net effect of these processes defines the dynamics of cells *T*_*i*_ in Eq. 2.

Doubly infected cells *T*_*ii*_ are produced when cells *T*_*i*_ are infected with virions *V*_*jh*_, following which reverse transcription again yields the provirus *i*. When a different provirus *j*(≠*i*) is produced, the result is the doubly infected cell *T*_*ij*_ carrying distinct proviruses *i* and *j*. Of course, cells *T*_*ij*_ can also be produced by the infections of cells *T*_*j*_ with another virion yielding the provirus *i*. Doubly infected cells too die with the rate constant *δ*. These processes are contained in Eqs. 3 and 4. We neglected cells infected more than twice, following experiments that suggest a rare occurrence of such cellular superinfection^90^.

Cells *T*_*i*_ and *T*_*ii*_ produce virions *V*_*ii*_ at the per capita rate *pξ*_*i*_, where *p* is the production rate of wild-type virions and *ξ*_*i*_ the production rate of virions *V*_*ii*_ relative the wild-type. *ξ*_*i*_ is thus the relative replicative fitness of genome *i*. Cells *T*_*hi*_ (*h*≠*i*) can also produce virions *V*_*ii*_. We assumed that viral RNA of the types *h* and *i* are present in cells *T*_*hi*_ in proportion to *ξ*_*h*_, and *ξ*_*i*_ respectively, and that they are randomly assorted into pairs and packaged into progeny virions. Thus, cells *T*_*hi*_ produce virions *V*_*ii*_ at the rate 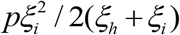 *V*_*hh*_ at the rate 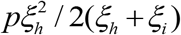 *V*_*ih*_ at the rate *pξ*_*h*_*ξ*_*i*_ / (*ξ*_*h*_ + *ξ*_*i*_). Virions are cleared at the rate *c*. These processes determine the dynamics of viral populations in Eqs. 5 and 6.

#### Mutation and recombination

We next describe the formalism to compute the probability *Q*_*i*_(*jh*). Following previous studies^49,50,91^, we let *R*_*k*_(*jh*) be the probability that genome *k* is produced by the recombination of genomes *j* and *h*, and *P_ik_* the probability that genome *k* mutates to genome *i*. Thus, *P*_*ik*_*R*_*k*_(*jh*) is the probability that genome *i* is produced from genomes *j* and *h* via the intermediate *k*. We recognize next that if genomes *j* and *h* differ in *d* positions, then recombination could produce a total of 2^*d*^ different genomes, depending on whether at each of the *d* positions, the nucleotide chosen is either from genome *j* or genome *h*. Summing over these different intermediates *k* yields the total probability of producing genome *i* from genomes *j* and *h* during reverse transcription:

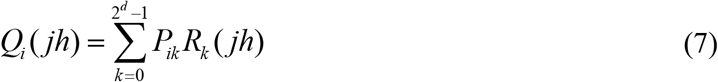

To compute *R*_*k*_(*jh*), we consider the desired path of the enzyme reverse transcriptase on the two genomes so that the enzyme is on the appropriate genome, *j* or *h*, at each of the *d* distinctive sites, so that the genome *k* is produced. We let the separation between the *x*^th^ and *x*+1^*st*^ distinctive sites be *l*_*x*_. If the enzyme has to be on the same genome (*j* or *h*) at both these sites, then it must perform an even number of crossovers in the length *l*_*x*_. Else, it must perform an odd number of crossovers. If the enzyme has a probability *p* of crossover per site, then the probabilities of even and odd crossovers over a length *l* are *P*_*even*_ (*l*) = (1+(1−2*ρ*)^*l*^)/2 and *P*_*odd*_ (*l*) = (1−(1−2*ρ*)^*l*^)/2, assuming that the crossover at any position is independent of the others and all crossovers happen with the same probability^91^. If we write *P*_*des*_(*x*+1|*x*) as the probability that the enzyme arrives on to the desired genome at the *x*+1*^st^* distinctive site given that it was on the desired genome at the *x*^*th*^ site, then depending on whether the associated crossovers must be even or odd, we write *P*_*des*_(*x*+1|*x*)=*P*_*even*_(*lx*) or *P*_*des*_(*x*+1|*x*)=*P_odd_*(*l*_*x*_). Note that *P*_*des*_(1)=1/2 because the enzyme could be on either genome at the start of the reverse transcription process. Thus, multiplying the probabilities over all the distinctive sites yields *R*_*k*_(*jh*):

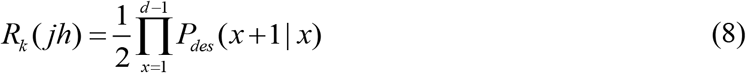

Next, we estimated the probability of producing genome *i* from genome *k*. If the two genomes differ at *u* sites, then genome *i* is produced from genome *k* by mutating genome *k* at the distinctive sites and nowhere else. Thus,

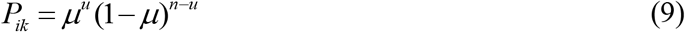

where *μ* is the per site mutation probability and *n* is the number of sites of interest.

#### Frequencies

Eqs. 1-9 yield a model of viral dynamics that predicts the growth of the populations of different mutants. We solved the equations for their steady states and obtained the corresponding frequencies of all proviruses, *ϕ*_*i*_, contained in productively infected cells:

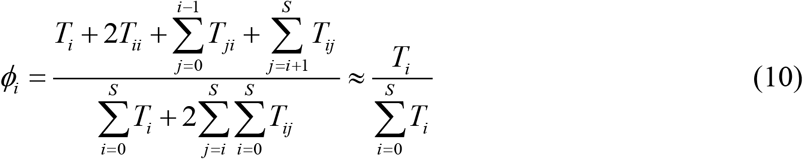

The approximation in Eq. 10 is justified by the small doubly infected cell population compared to the singly infected cell population. The parameter values employed are in Table 1. *ϕ*_*i*_ yield the frequencies of various VRC01-resistant mutants before the start of treatment. Previous studies have shown that this deterministic formalism yields mutant frequencies in agreement with stochastic population genetics-based simulations of HIV evolution^50^. The frequencies are expected to hold also for the proviruses in latently infected cells. Further, we expect ART not to influence the latter frequencies; standard first-line ART drugs target HIV reverse transcriptase and protease, whereas VRC01 targets the HIV envelope. We assumed therefore that following ART, the frequencies of VRC01-resistant strains contained in the latent reservoir are given by Eq. 10.

We focused next on the reactivation of latently infected cells carrying the resistant genomes.

### Latency reactivation and viral rebound

We developed a stochastic framework to describe the reactivation of latently infected cells carrying mutant proviruses resistant to VRC01. Because reactivation is most likely of the cells carrying the fittest mutant, which are the most prevalent, we considered cells *L*_*m*_ carrying the fittest mutant strain above. Previous studies on ART resistance too have considered the dominant mutant alone.^53^ The reactivation of these cells and the ensuing growth of mutant virions is then described by the following events:

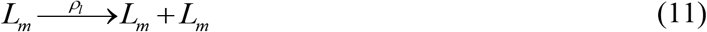

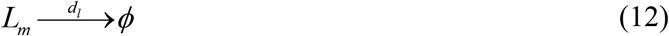

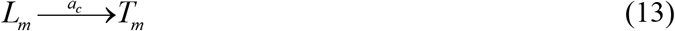

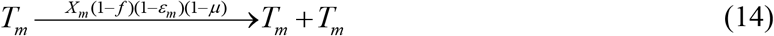

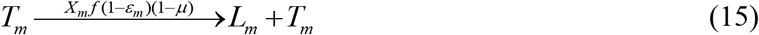

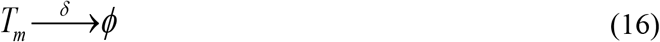

Here, cells *L*_*m*_ proliferate at the per capita rate *ρ*_*l*_ (Eq. 11), die at the per capita rate *d*_*l*_ (Eq. 12), and get activated to productively infected cells, *T*_*m*_, at the per capita rate *a*_*c*_ (Eq. 13). Cells *T*_*m*_ carry the mutant provirus and produce resistant virions, *V*_*mm*_, at the per capita rate *pξ*_*m*_, which in turn infect uninfected cells, *U*, with the rate constant *kβ*_*mm*_. Here, *ξ*_*mm*_ and *β_mm_* are the relative replicative ability and infectivity, respectively, of the strain *m*. Free virions are cleared at the per capita rate *c*. Viral production and clearance are typically rapid^92^ compared to changes in cell populations, so that following an approximation widely used (e.g., see^48^), *V*_*mm*_ can be assumed to be in pseudo-state with *T*_*m*_. Thus, *V*_*mm*_≈*pξ*_*m*_*T*_*m*_/*c*. The rate, *kβ*_*mm*_*VU*, of the infection of *U* thus becomes *kβ*_*mm*_*pξmT*_*m*_*U/c*. We recognized that *U* is not altered significantly due to new infections, especially following ART when the viremia is small. We let a fraction *f* of the new infections lead to latency and the remaining to productive infection. Further, we let a VRC01-resistant strain mutate back to the wild-type with a probability *μ*, the point mutation rate of HIV-1. The resulting description would capture scenarios where a single mutation is adequate to develop resistance, as is the case with VRC01^27^. Thus, the rate *kβ*_*mm*_*pξ*_*m*_*T*_*m*_*U*(1−*f*)(1−*μ*)/*c* becomes the rate of the growth of *T*_*m*_ in the absence of therapy. If VRC01 were to block infections with efficacy *ε*_*m*_, the rate would become *X*_*m*_*T*_*m*_(1−*f*)(1−*μ*)(1−*ε*_*m*_)*/c* with *X*_*m*_=*kβ*_*mm*_*pξ*_*m*_*U/c*. Thus, we let *T*_*m*_ (effectively) double with the per capita rate *X*_*m*_(1−*f*)(1−*μ*)(1−*ε*_*m*_)/*c* (Eq. 14), and yield new latently infected cells at the per capita rate *X*_*m*_*f*(1−*μ*)(1−*ε*_*m*_)/*c* (Eq. 15). Cells *T*_*m*_ die with the rate constant *δ*. These events together describe the growth of resistance to VRC01 and therapy failure.

We solved the model equations using the Gillespie algorithm^93^ with parameter values representative of HIV-1 infection in the presence of VRC01 therapy. The model was implemented using a program written in MATLAB. The parameter estimates are listed in Tables 1 and 2. The initial population of *L*_*m*_ was set by the frequencies of mutants estimated. With each parameter setting, we performed 500 realizations to examine the dynamics of treatment failure.

### Comparison with clinical data

We applied our model to predict the outcomes of trials with the VRC01^17^. For this, we explicitly considered VRC01 pharmacokinetics. Further, we created a virtual patient population to account for the inter-patient variations seen in the trials. In each patient, we performed stochastic simulations and identified the VRC01 failure time following the model above. We compared the resulting distribution of failure times with those observed in the trials. We describe the methods we used here.

#### Data

We considered data of viral resurgence following VRC01 therapy during ATIs from two clinical trials, the A5340 trial and the NIH trial^17^. In the A5340 trial, 14 adult chronic HIV patients with a median ART duration of 4.7 years and viremia maintained below detection were administered 3 doses of VRC01 (40 mg/kg of body weight) intravenously, starting a week before the cessation of ART and with 3 week intervals. Plasma viremia was measured weekly to check for rebound (>20 copies/ml). Of the 14 participants, 1 participant stopped ART before VRC01 administration and was not part of the trial data analysis. Further, 1 participant maintained remission until well after the 3 doses were administered and thus appeared not to have suffered failure from VRC01 resistance. We considered data from the remaining 12 patients, who experienced virological breakthrough between 2 and 8 weeks from the discontinuation of ART. For each patient, we assumed the breakthrough time was anywhere within the week from the last undetectable to the first detectable viral load measurement.

The NIH trial had 10 participants with similar characteristics as the A5340 trial. The participants had been on ART much longer, however, with a median duration of 10 years, before entry into the trial. ART was stopped 3 days after the first VRC01 administration. VRC01 doses, at the same dosage as above, were administered subsequently on days 14 and 28 post ART cessation and once every month thereafter. Plasma viremia was measured every week for 1 month and then every 2 weeks until 6 months. The patients experienced virological breakthrough between 2 and 12 weeks from the discontinuation of ART. We considered this time of virological failure with uncertainties as defined above.

We note that both trials did not screen for pre-existing VRC01 resistance. While the primary endpoints of the trials were safety and tolerance, secondary endpoints were viral remission, based on which the trial data report Kaplan-Meier survival curves^17^. We considered the latter data too in our analysis.

#### VRC01 pharmacokinetics and pharmacodynamics

We let the efficacy of VRC01 against the drug-resistant mutant strain, *ε*_*m*_, be related to its plasma concentration, *A*, by the Hill function^49,55,94^,

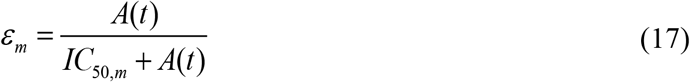

where *IC*_*50,m*_ is the value of *A* at which VRC01 is 50% efficacious. For simplicity, we set the Hill coefficient to 1. The antibody concentration has been observed to decline in a biphasic manner upon dosing^25^. We therefore described the time course of the antibody concentration using the following expression when successive doses are administered at time points *ω*_1_, *ω*_2_ and so on:

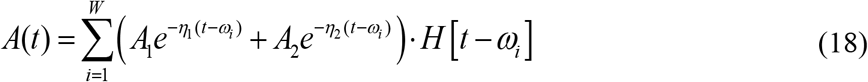

Here, *A*_1_ and *A*_2_ are the portions of a dose that decay in the first and second phases, respectively, with decay rates *η*_1_, and *η*_2_ and *W* is the total number of doses. We estimated the pharmacokinetic parameters using fits of the above equation to data of VRC01 concentrations following a single dose^25^.

#### ART efficacy

For the duration that ART is used simultaneously with VRC01, we replaced the term (1−*ε*_*m*_) in Eqs. 14 and 15 by 1 − *ε*_*comb*_ = 1 − *ε*_*m*_(1– *ε*_*ART*_), where *ε*_*ART*_ is the efficacy of ART and *ε*_*comb*_ is the combined efficacy of VRC01 and ART. We let *ε*_*ART*_ be constant while on ART and set it to zero thereafter.

#### Virtual patient population

To capture inter-patient variations in the response to bNAb therapy, we constructed a virtual population of clinical trial participants as follows. For simplicity, we considered variations in two factors, the initial latent pool size, *L*_0_, and the viral production rate, *p*, across individuals.

We assumed that variations in all host factors could be subsumed in the variation in *L*_0_ and that variations in viral factors could be subsumed in *p*. We let *L*_0_ vary log-normally and *p* normally across individuals. Thus, the corresponding density functions were

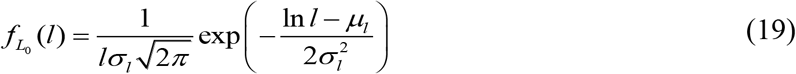

and

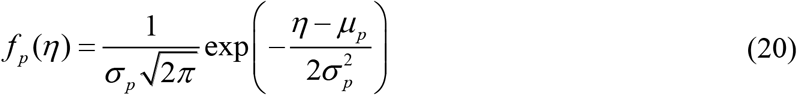

where *μ*_*l*_ and *σ*_*l*_ are the mean and standard deviation of *lnL*_0_ and *μ*_*p*_ and *σ*_*p*_ are the mean and standard deviation of *p*, respectively. The parameters in the distributions were chosen based on earlier studies or to fit the results of the A5340 trial. From the resulting distributions, we sampled many pairs (≥ 10000) of values of *L*_0_ and *p*, with each pair representing an individual. We thus created a virtual patient population, which we then subjected to VRC01 therapy according to the protocols in the respective trials. For each individual, we ran a stochastic simulation (Eqs. 1–18) and examined the dynamics of virological breakthrough. From the resulting dynamics, we constructed the distribution of breakthrough times and used it to build Kaplan-Meier survival plots sampling virtual patients mimicking the clinical trial protocols.

## ACKNOWLEDGMENTS

This work was supported by the Wellcome Trust/DBT India Alliance Senior Fellowship IA/S/14/1/501307 (NMD).

## REFERENCES

1. Arts, E. J. & Hazuda, D. J. HIV-1 antiretroviral drug therapy. Cold Spring Harb. Perspect. Med. 2, a007161–a007161 (2012).

2. Chun, T. W., Davey, R. T., Engel, D., Lane, H. C. & Fauci, A. S. AIDS: Re-emergence of HIV after stopping therapy. Nature 401, 874–875 (1999).

3. Chun, T. W., Moir, S. & Fauci, A. S. HIV reservoirs as obstacles and opportunities for an HIV cure. Nat. Immunol. 16, 584–589 (2015).

4. Chun, T. W. et al. Early establishment of a pool of latently infected, resting CD4+ T cells during primary HIV-1 infection. Proc. Natl. Acad. Sci. U. S. A. 95, 8869–8873 (1998).

5. Hosmane, N. N. et al. Proliferation of latently infected CD4+ T cells carrying replication-competent HIV-1: Potential role in latent reservoir dynamics. J. Exp. Med. 214, 959–972 (2017).

6. Reeves, D. B. et al. A majority of HIV persistence during antiretroviral therapy is due to infected cell proliferation. Nat. Commun. 9, 4811 (2018).

7. Finzi, D. et al. Latent infection of CD4+ T cells provides a mechanism for lifelong persistence of HIV-1, even in patients on effective combination therapy. Nat. Med. 5, 512–517 (1999).

8. Weinberger, L. S., Burnett, J. C., Toettcher, J. E., Arkin, A. P. & Schaffer, D. V. Stochastic gene expression in a lentiviral positive-feedback loop: HIV-1 Tat fluctuations drive phenotypic diversity. Cell 122, 169–182 (2005).

9. Ho, Y. C. et al. Replication-competent noninduced proviruses in the latent reservoir increase barrier to HIV-1 cure. Cell 155, 540–551 (2013).

10. Sáez-Cirión, A. et al. Post-treatment HIV-1 controllers with a long-term virological remission after the interruption of early initiated antiretroviral therapy ANRS VISCONTI study. PLoS Pathog. 9, e1003211 (2013).

11. Deeks, S. G. et al. International AIDS Society global scientific strategy: Towards an HIV cure 2016. Nat. Med. 22, 839–850 (2016).

12. Cockerham, L. R., Hatano, H. & Deeks, S. G. Post-treatment controllers: Role in HIV “cure” research. Curr. HIV/AIDS Rep. 13, 1–9 (2016).

13. Halper-Stromberg, A. & Nussenzweig, M. C. Towards HIV-1 remission: Potential roles for broadly neutralizing antibodies. J. Clin. Invest. 126, 415–423 (2016).

14. Wen, Y., Bar, K. J. & Li, J. Z. Lessons learned from HIV antiretroviral treatment interruption trials. Curr. Opin. HIV AIDS 13, 416–421 (2018).

15. Zhou, T. et al. Structural basis for broad and potent neutralization of HIV-1 by antibody VRC01. Science 329, 811–817 (2010).

16. Wu, X. et al. Rational design of envelope identifies broadly neutralizing human monoclonal antibodies to HIV-1. Science 329, 856–861 (2010).

17. Bar, K. J. et al. Effect of HIV antibody VRC01 on viral rebound after treatment interruption. N. Engl. J. Med. 375, 2037–2050 (2016).

18. Caskey, M. et al. Viraemia suppressed in HIV-1-infected humans by broadly neutralizing antibody 3BNC117. Nature 522, 487–491 (2015).

19. Lu, L. L., Suscovich, T. J., Fortune, S. M. & Alter, G. Beyond binding: Antibody effector functions in infectious diseases. Nat. Rev. Immunol. 18, 46–61 (2018).

20. Nishimura, Y. et al. Early antibody therapy can induce long-lasting immunity to SHIV. Nature 543, 559–563 (2017).

21. Borducchi, E. N. et al. Antibody and TLR7 agonist delay viral rebound in SHIV-infected monkeys. Nature 563, 360–364 (2018).

22. Rudicell, R. S. et al. Enhanced potency of a broadly neutralizing HIV-1 antibody in vitro improves protection against lentiviral infection in vivo. J. Virol. 88, 12669–12682 (2014).

23. Li, J. Z. et al. The size of the expressed HIV reservoir predicts timing of viral rebound after treatment interruption. AIDS 30, 343–353 (2016).

24. Morris, L. & Mkhize, N. N. Prospects for passive immunity to prevent HIV infection. PLoS Med. 14, e1002436 (2017).

25. Lynch, R. M. et al. Virologic effects of broadly neutralizing antibody VRC01 administration during chronic HIV-1 infection. Sci. Transl. Med. 7, 319ra206 (2015).

26. Mayer, K. H. et al. Safety, pharmacokinetics, and immunological activities of multiple intravenous or subcutaneous doses of an anti-HIV monoclonal antibody, VRC01, administered to HIV-uninfected adults: Results of a phase 1 randomized trial. PLoS Med. 14, e1002435 (2017).

27. Lynch, R. M. et al. HIV-1 fitness cost associated with escape from the VRC01 class of CD4 binding site neutralizing antibodies. J. Virol. 89, 4201–4213 (2015).

28. Otsuka, Y. et al. Diverse pathways of escape from all well-characterized VRC01-class broadly neutralizing HIV-1 antibodies. PLoS Pathog. 14, e1007238 (2018).

29. Tachibana, S. et al. A 2-4-amino acid deletion in the V5 region of HIV-1 env gp120 confers viral resistance to the broadly neutralizing human monoclonal antibody, VRC01. AIDS Res. Hum. Retroviruses 33, 1248–1257 (2017).

30. Wang, W. et al. N463 glycosylation site on V5 loop of a mutant gp120 regulates the sensitivity of HIV-1 to neutralizing monoclonal antibodies VRC01/03. J. Acquir. Immune Defic. Syndr. 69, 270–277 (2015).

31. Van Laethem, K., Theys, K. & Vandamme, A. M. HIV-1 genotypic drug resistance testing: Digging deep, reaching wide? Curr. Opin. Virol. 14, 16–23 (2015).

32. Bruner, K. M. et al. A quantitative approach for measuring the reservoir of latent HIV-1 proviruses. Nature 566, 120–125 (2019).

33. Silver, N. et al. Characterization of minority HIV-1 drug resistant variants in the United Kingdom following the verification of a deep sequencing-based HIV-1 genotyping and tropism assay. AIDS Res. Ther. 15, 18 (2018).

34. Estes, J. D. et al. Defining total-body AIDS-virus burden with implications for curative strategies. Nat. Med. 23, 1271–1276 (2017).

35. Conway, J. M. & Coombs, D. A stochastic model of latently infected cell reactivation and viral blip generation in treated HIV patients. PLoS Comput. Biol. 7, 1–25 (2011).

36. Hill, A. L. Modeling HIV persistence and cure studies. Curr. Opin. HIV AIDS 13, 428–434 (2018).

37. Conway, J. M. & Perelson, A. S. Post-treatment control of HIV infection. Proc. Natl. Acad. Sci. U. S. A. 6, 5467–5472 (2015).

38. Dar, R. D., Hosmane, N. N., Arkin, M. R., Siliciano, R. F. & Weinberger, L. S. Screening for noise in gene expression identifies drug synergies. Science 344, 1392–1396 (2014).

39. Hill, A. L., Rosenbloom, D. I. S., Fu, F., Nowak, M. A. & Siliciano, R. F. Predicting the outcomes of treatment to eradicate the latent reservoir for HIV-1. Proc. Natl. Acad. Sci. U. S. A. 111, 13475–13480 (2014).

40. Rong, L. & Perelson, A. S. Modeling latently infected cell activation: Viral and latent reservoir persistence, and viral blips in HIV-infected patients on potent therapy. PLoS Comput. Biol. 5, e1000533 (2009).

41. Conway, J. M., Perelson, A. S. & Li, J. Z. Predictions of time to HIV viral rebound following ART suspension that incorporate personal biomarkers. PLoS Comput. Biol. 15, e1007229 (2019).

42. Pinkevych, M. et al. HIV reactivation from latency after treatment interruption occurs on average every 5-8 days—implications for HIV remission. PLoS Pathog. 11, e1005000 (2015).

43. Gupta, V. & Dixit, N. M. Trade-off between synergy and efficacy in combinations of HIV-1 latency-reversing agents. PLoS Comput. Biol. 14, e1006004 (2018).

44. Althaus, C. L. & De Boer, R. J. Intracellular transactivation of HIV can account for the decelerating decay of virus load during drug therapy. Mol. Syst. Biol. 6, 1–8 (2010).

45. Rouzine, I. M., Weinberger, A. D. & Weinberger, L. S. An evolutionary role for HIV latency in enhancing viral transmission. Cell 160, 1002–1012 (2015).

46. Davenport, M. P. et al. Functional cure of HIV: the scale of the challenge. Nat. Rev. Immunol. 19, 45–54 (2019).

47. Ribeiro, R. M., Bonhoeffer, S. & Nowak, M. A. The frequency of resistant mutant virus before antiviral therapy. AIDS 12, 461–465 (1998).

48. Ribeiro, R. M. & Bonhoeffer, S. Production of resistant HIV mutants during antiretroviral therapy. Proc. Natl. Acad. Sci. U. S. A. 97, 7681–7686 (2000).

49. Arora, P. & Dixit, N. M. Timing the emergence of resistance to anti-HIV drugs with large genetic barriers. PLoS Comput. Biol. 5, e1000305 (2009).

50. Gadhamsetty, S. & Dixit, N. M. Estimating frequencies of minority nevirapine-resistant strains in chronically HIV-1-infected individuals naive to nevirapine by using stochastic simulations and a mathematical model. J. Virol. 84, 10230–10240 (2010).

51. Pennings, P. S. Standing genetic variation and the evolution of drug resistance in HIV. PLoS Comput. Biol. 8, e1002527 (2012).

52. Tripathi, K., Balagam, R., Vishnoi, N. K. & Dixit, N. M. Stochastic simulations suggest that HIV-1 survives close to its error threshold. PLoS Comput. Biol. 8, e1002684 (2012).

53. Rosenbloom, D. I. S., Hill, A. L., Rabi, S. A., Siliciano, R. F. & Nowak, M. A. Antiretroviral dynamics determines HIV evolution and predicts therapy outcome. Nat. Med. 18, 1378–1385. (2012).

54. Hinkley, T. et al. A systems analysis of mutational effects in HIV-1 protease and reverse transcriptase. Nat. Genet. 43, 487–489 (2011).

55. Brandenberg, O. F. et al. Predicting HIV-1 transmission and antibody neutralization efficacy in vivo from stoichiometric parameters. PLoS Pathog. 13, e1006313 (2017).

56. Shet, A., Nagaraja, P. & Dixit, N. M. Viral Decay Dynamics and Mathematical Modeling of Treatment Response. J. Acquir. Immune Defic. Syndr. 73, 245–251 (2016).

57. Maree, A. F. M., Keulen, W., Boucher, C. A. B. & De Boer, R. J. Estimating relative fitness in viral competition experiments. J. Virol. 74, 11067–11072 (2000).

58. Salantes, D. B. et al. HIV-1 latent reservoir size and diversity are stable following brief treatment interruption. J. Clin. Invest. 128, 3102–3115 (2018).

59. Kepler, T. B. & Perelson, A. S. Drug concentration heterogeneity facilitates the evolution of drug resistance. Proc. Natl. Acad. Sci. U. S. A. 95, 11514–11519 (1998).

60. Goulder, P. & Deeks, S. G. HIV control: Is getting there the same as staying there? PLoS Pathog. 14, e1007222 (2018).

61. Fraser, C. et al. Virulence and pathogenesis of HIV-1 infection: An evolutionary perspective. Science 343, 1243727 (2014).

62. Prague, M., Commenges, D., Guedj, J., Drylewicz, J. & Thiébaut, R. NIMROD: A program for inference via a normal approximation of the posterior in models with random effects based on ordinary differential equations. Comput. Methods Programs Biomed. 111, 447–458 (2013).

63. Lavielle, M. Mixed Effects Models for the Population Approach: Models, Tasks, Methods and Tools. Chapman and Hall/CRC (2014).

64. de Bree, G. J. & Sanders, R. W. Broadly neutralising antibodies in post-treatment control. Lancet HIV 6, e271–e272 (2019).

65. Haynes, B. F., Burton, D. R. & Mascola, J. R. Multiple roles for HIV broadly neutralizing antibodies. Sci. Transl. Med. 11, eaaz2686 (2019).

66. Schommers, P. et al. Restriction of HIV-1 escape by a highly broad and potent neutralizing antibody. Cell 180, 471–489.e22 (2020).

67. Keele, B. F. et al. Identification and characterization of transmitted and early founder virus envelopes in primary HIV-1 infection. Proc. Natl. Acad. Sci. U. S. A. 105, 7552–7557 (2008).

68. Shankarappa, R. et al. Consistent viral evolutionary changes associated with the progression of human immunodeficiency virus type 1 infection. J. Virol. 73, 10489–10502 (1999).

69. Crowell, T. A. et al. Safety and efficacy of VRC01 broadly neutralising antibodies in adults with acutely treated HIV (RV397): a phase 2, randomised, double-blind, placebo-controlled trial. Lancet HIV 6, e297–e306 (2019).

70. Hashimoto, M. et al. CD8 T cell exhaustion in chronic infection and cancer: Opportunities for interventions. Annu. Rev. Med. 69, 301–318 (2018).

71. McLane, L. M., Abdel-Hakeem, M. S. & Wherry, E. J. CD8 T cell exhaustion during chronic viral infection and cancer. Annu. Rev. Immunol. 37, 457–495(2019).

72. Johnson, P. L. F. et al. Vaccination alters the balance between protective immunity, exhaustion, escape, and death in chronic infections. J. Virol. 85, 5565–5570 (2011).

73. Baral, S., Roy, R. & Dixit, N. M. Modeling how reversal of immune exhaustion elicits cure of chronic hepatitis C after the end of treatment with direct-acting antiviral agents. Immunol. Cell Biol. 96, 969–980 (2018).

74. Baral, S., Antia, R. & Dixit, N. M. A dynamical motif comprising the interactions between antigens and CD8 T cells may underlie the outcomes of viral infections. Proc. Natl. Acad. Sci. U. S. A. 116, 17393–17398 (2019).

75. Baral, S., Raja, R., Sen, P. & Dixit, N. M. Towards multiscale modeling of the CD8+ T cell response to viral infections. Wiley Interdiscip Rev Syst Biol Med 11, e1446 (2019).

76. Wu, X. et al. Focused evolution of HIV-1 neutralizing antibodies revealed by structures and deep sequencing. Science 333, 1593–1602 (2011).

77. Wu, X. et al. Maturation and diversity of the VRC01-antibody lineage over 15 years of chronic HIV-1 infection. Cell 161, 470–485 (2015).

78. Scheid, J. F. et al. Sequence and structural convergence of broad and potent HIV antibodies that mimic CD4 binding. Science 333, 1633–1637 (2011).

79. Gilbert, P. B. et al. Basis and statistical design of the passive HIV-1 antibody mediated prevention (AMP) test-of-concept efficacy trials. Stat. Commun. Infect. Dis. 9, 20160001 (2017).

80. Huang, Y. et al. Modeling cumulative overall prevention efficacy for the VRC01 phase 2b efficacy trials. Hum. Vaccines Immunother. 14, 2116–2127 (2018).

81. Raja, R., Pareek, A., Newar, K. & Dixit, N. M. Mutational pathway maps and founder effects define the within-host spectrum of hepatitis C virus mutants resistant to drugs. PLoS Pathog. 15, e1007701 (2019).

82. Prague, M. et al. Viral rebound kinetics following single and combination immunotherapy for HIV / SIV. bioRxiv 1–72 (2019) doi:10.1101/700401.

83. Cardozo, E. F. et al. Treatment with integrase inhibitor suggests a new interpretation of HIV RNA decay curves that reveals a subset of cells with slow integration. PLoS Pathog. 13, e1006478 (2017).

84. Schoofs, T. et al. HIV-1 therapy with monoclonal antibody 3BNC117 elicits host immune responses against HIV-1. Science 352, 997–1001 (2016).

85. Garg, A. K., Desikan, R. & Dixit, N. M. Preferential presentation of high-affinity immune complexes in germinal centers can explain how passive immunization improves the humoral response. Cell Rep. 29, 3946–3957.e5 (2019).

86. DiLillo, D. J. & Ravetch, J. V. Differential Fc-receptor engagement drives an anti-tumor vaccinal effect. Cell 161, 1035–1045 (2015).

87. Desikan, R., Raja, R. & Dixit, N. M. Early exposure to broadly neutralizing antibodies may trigger a dynamical switch from progressive disease to lasting control of SHIV infection. bioRxiv 548727 (2020) doi:10.1101/548727.

88. Chen, B. K., Gandhi, R. T. & Baltimore, D. CD4 down-modulation during infection of human T cells with human immunodeficiency virus type 1 involves independent activities of vpu, env, and nef. J. Virol. 70, 6044–6053 (1996).

89. Dixit, N. M. & Perelson, A. S. HIV dynamics with multiple infections of target cells. Proc. Natl. Acad. Sci. U. S. A. 102, 8198–8203 (2005).

90. Josefsson, L. et al. Majority of CD4 + T cells from peripheral blood of HIV-1-infected individuals contain only one HIV DNA molecule. Proc. Natl. Acad. Sci. U. S. A. 108, 11199–11204 (2011).

91. Nagaraja, P., Alexander, H. K., Bonhoeffer, S. & Dixit, N. M. Influence of recombination on acquisition and reversion of immune escape and compensatory mutations in HIV-1. Epidemics 14, 11–25 (2016).

92. Ramratnam, B. et al. Rapid production and clearance of HIV-1 and hepatitis C virus assessed by large volume plasma apheresis. Lancet 354, 1782–1785 (1999).

93. Gillespie, D. T. Exact stochastic simulation of coupled chemical reactions. J. Phys. Chem. 81, 2340–2361 (1977).

94. Dixit, N. M. & Perelson, A. S. Complex patterns of viral load decay under antiretroviral therapy: influence of pharmacokinetics and intracellular delay. J. Theor. Biol. 226, 95–109 (2004).

