## Supplementary material for "Pre-existing resistant proviruses can compromise maintenance of remission by VRC01 in chronic HIV-1 infection"

### for

#### Text S1. VRC01 therapy failure is unlikely to be due to *de novo* mutation

Here, we examined the alternative possibility of the failure of VRC01 therapy due to the reactivation of latently infected cells carrying VRC01-sensitive strains followed by mutation-driven development of resistance. We therefore considered latently infected cells carrying wild-type or VRC01-sensitive genomes alone. For therapy failure, such cells would have to become activated, produce virions, which would have to escape VRC01 and immune pressure, infect other cells, and in the process mutate to yield VRC01-resistant genomes. The latter genomes would have to escape immune pressure too and establish progressive infection. We constructed first a stochastic model to describe these processes.

**Stochastic model.** The following events captured the processes leading to the development of VRC01 resistance.

$$L \xrightarrow{\rho_l} L + L \tag{S1}$$

$$L \xrightarrow{a_c} T \tag{S2}$$

$$L \xrightarrow{d_l} \phi \tag{S3}$$

$$T \xrightarrow{X(1-f)(1-\varepsilon)(1-\mu)} T + T \tag{S4}$$

$$T \xrightarrow{Xf(1-\varepsilon)(1-\mu)} L + T \tag{S5}$$

$$T \xrightarrow{X(1-f)(1-\varepsilon)\mu} T_m + T \tag{S6}$$

$$T \xrightarrow{Xf(1-\varepsilon)\mu} L_m + T \tag{S7}$$

$$T \xrightarrow{\delta} \phi \tag{S8}$$

$$L_m \xrightarrow{\rho_l} L_m + L_m \tag{S9}$$

$$L_m \xrightarrow{a_c} T_m \tag{S10}$$

$$L_m \xrightarrow{d_l} \phi \tag{S11}$$

Here, latently infected cells, $L$, proliferate at the per capita rate $\rho_l$ (Eq. S1), get activated to productively infected cells, $T$, at the per capita rate $a_c$ (Eq. S2), and die at the per capita rate $d_l$ (Eq. S3). Cells $T$ produce virions, $V$, at the per capita rate $p$, which in turn infect uninfected cells, $U$, with the rate constant $k$. Free virions are cleared at the per capita rate $c$. Viral production and clearance are typically rapid[1] compared to changes in cell populations, so that following previous studies, $V$ can be assumed to be in pseudo-state with $T$. Thus, $V \approx pT/c$. The rate, $kVU$, of the infection of $U$ thus becomes $kpTU/c$. We recognized that $U$ is not altered significantly due to new infections, especially following ART when the viremia is small. We let a fraction $f$ of the new infections lead to latency and the remaining to productive infection. Further, we let a VRC01-resistant strain be produced during reverse transcription of a wild-type viral genome with a probability $\mu$, the point mutation rate of HIV-1. The resulting description would capture scenarios where a single mutation is adequate to develop high level resistance to a bNAb, as is the case with VRC01[2]. Thus, the rate $kpTU(1\text{-}f)(1\text{-}\mu)/c$ becomes the rate of the growth of $T$ in the absence of therapy. If VRC01 blocked infections with efficacy $\varepsilon$, the rate would become $XT(1\text{-}f)(1\text{-}\mu)(1\text{-}\varepsilon)/c$ with $X=kpU/c$. Thus, we let $T$ (effectively) double with the per capita rate $X(1\text{-}f)(1\text{-}\mu)(1\text{-}\varepsilon)/c$ (Eq. S4), and yield new latently infected cells at the per capita rate $Xf(1\text{-}\mu)(1\text{-}\varepsilon)/c$ (Eq. S5). Similarly, the infections would lead to productively infected cells, $T_m$, and latently infected cells, $L_m$, carrying proviruses resistant to VRC01 at the per capita rates $X(1\text{-}f)\mu(1\text{-}\varepsilon)/c$ (Eq. S6) and $Xf\mu(1\text{-}\varepsilon)/c$ (Eq. S7), respectively. Cells $T$ die at the per capita rate $\delta$ (Eq. S8). We let cells $L_m$ also proliferate at the per capita rate $\rho_l$ (Eq. S9), get activated to $T_m$ at the per capita rate $a_c$ (Eq. S10), and die at the per capita rate $d_l$ (Eq. S11), akin to cells $L$. Our interest is in the formation of the first cell $T_m$, at which point, we assume that resistance to the bNAb has arisen *de novo*. We do recognize that detectable viral rebound requires the cell $T_m$ to produce virus before it dies and that the resulting viruses escape drug and immune pressure and establish sustained infection. Our simulations can readily be extended to account for the latter events, as has also been done in other branching process formulations[3,4]. The time for the first formation of $T_m$ may thus be viewed as a lower bound on the time for establishing successful infection.

We solved the model equations using the Gillespie algorithm[5] with parameter values representative of HIV-1 infection in the presence of VRC01 therapy (Tables 1 and 2). The model was implemented using a program written in MATLAB. We integrated the model until 21 d, comparable to failure times seen with VRC01 therapy in clinical trials[6]. Averaging over 500 realizations, we estimated the mean waiting time for the emergence of $T_m$.

When the mean waiting time was much larger than 21 d and the initial latent pool was large, stochastic simulations became prohibitive. We therefore constructed a deterministic model to estimate the mean waiting time, which not only provided a test of our simulations, but also allowed estimation of the waiting times beyond the regimes where stochastic simulations were feasible.

**Deterministic waiting time model.** In a previous study, a deterministic multi-strain model was constructed to estimate the mean waiting time for the emergence of resistance to antiretroviral drugs that offered large genetic barriers to resistance[7]. The model recapitulated the failure of such drugs *in vitro* and has since been extended to describe the accumulation of immune escape mutations[8]. Latently infected cells were not central to drug resistance or immune escape and therefore were not considered in the model. Here, we adapted the model to describe the failure of VRC01 therapy via the activation of latently infected cells and the subsequent occurrence of mutations. Following the arguments above, we let a single mutation lead to VRC01 resistance. The resulting equations were as follows.

$$\frac{dL}{dt} = (\rho - d_l)L + kUVf(1-\mu)(1-\varepsilon) - H[t-\tau]\cdot a_c L \quad \text{(S12)}$$

$$\frac{dT}{dt} = H[t-\tau]\cdot a_c L + kUV(1-f)(1-\mu)(1-\varepsilon) - \delta T \quad \text{(S13)}$$

$$\frac{dV}{dt} = pT - cV \quad \text{(S14)}$$

$$\frac{dT_m}{dt} = H[t-\tau_m]\cdot kUV(1-f)\mu(1-\varepsilon) - \delta T_m \quad \text{(S15)}$$

The variables and parameters in the equations have the same meanings as those in the model above (Eqs. S1-S11). The formalism uses expressions for the growth and decay of the various cells and virions that follow standard models of viral dynamics[9,10]. Further, it accounts for the delay in the occurrence of rare events by estimating their mean waiting times. Thus, here, $\tau$ and $\tau_m$ represent the mean waiting times for the formation of the first productively infected cell carrying a VRC01-sensitive strain and the first productively infected cell carrying a VRC01-resistant strain, respectively. The model considered dynamics following the successful control of viremia by ART, so that initially latently infected cells alone existed. Further, here, the cells were all assumed to carry sensitive viral genomes. The dynamics of these cells, $L$, was determined by their proliferation and death rates (Eq. S1). They could also be activated stochastically to productively infected cells, $T$. To capture this stochastic effect, we assumed that the first such reactivation event occurred at time $\tau$ (estimated below). Until this time, $T$=0 and no cells $L$ were lost due to reactivation. At $t=\tau$, we set $T$=1 and let the reactivation of cells $L$ proceed at the per capita rate $a_c$. We implemented this waiting time using the Heaviside function, defined as $H[t-\tau]=0$ when $t<\tau$ and $H[t-\tau]=1$ otherwise (Eqs. S12 and S13). Because each productively infected cell produces many virions, which in turn can infect uninfected cells and expand $T$ at a much faster rate than the reactivation of $L$, we let the subsequent dynamics of $T$ proceed deterministically (Eq. S13). We considered virus production from $T$ and viral clearance explicitly (Eq. S14). Note that cells $T$ can only produce sensitive virions. When these virions infect target cells, they can give rise to cells $T_m$, productively infected with resistant mutants. Again, because mutations are intrinsically rare, we let the first cell $T_m$ be formed at its mean waiting time, $\tau_m$, implemented using the Heaviside function, $H(t-\tau_m)$ (Eq. S15). We recognized that $T_m$ could be formed by another pathway: Infection with a wild-type virion could lead to a latently infected cell carrying a resistant provirus because of mutation during reverse transcription. Reactivation of the latter latently infected cell could yield $T_m$. This pathway, which is accounted for in the stochastic model above, is far less likely than the formation of $T_m$ by infection of uninfected cells and we therefore ignored it in our deterministic model. The close agreement between the deterministic and stochastic models, shown below, justified this approximation.

To estimate $\tau$, we recognized that the time, $w$, to the emergence of the first cell $T$ can be modelled using a time-inhomogeneous Poisson process with rate $a_c L(t)$. The probability that $w$ is smaller than $s \in [0,\infty)$ is then $P(w<s) = 1-\exp(-a_c \int_0^s L(t)dt)$. The mean waiting time is thus

$\tau = \langle w \rangle = \int_0^\infty s(\partial P / \partial s)ds$ , or

$$\tau = -\int_0^\infty s \frac{\partial}{\partial s}\left( \exp\left( -a_c \int_0^s L(t)dt \right)\right) ds \qquad \text{(S16)}$$

Similarly, if we defined $y$ as the time to the emergence of the first cell $T_m$, then from Eq. S15, assuming $U$ does not vary significantly with time, it followed that $P(y<s) = 1-\exp(-kU(1-f)\mu(1-\varepsilon)\int_0^s V(t)dt)$, which in turn yielded

$$\tau_m = -\int_0^\infty s \frac{\partial}{\partial s}\left( \exp\left( -kU(1-f)\mu(1-\varepsilon)\int_0^s V(t)dt \right)\right) ds \qquad \text{(S17)}$$

Note that $\partial P(y<s)/\partial s = f_y(s)$ is the probability density function of the waiting time *y*. We solved Eqs. S12-S17 using a numerical algorithm developed earlier[7]. The evolution of the quantities *L*, *T*, *V* and $T_m$ depend on estimates of $\tau$ and $\tau_m$ (Eqs. S12-S15), which in turn depend on estimates of *L* and *V* (Eqs. S16 and S17). This coupling of the equations makes them difficult to solve. The algorithm decouples the equations by obtaining approximate estimates of $\tau$ and $\tau_m$ at each time step in the numerical integration of the differential equations (Eqs. S12-S15) based on current sizes of *L* and *V*. The parameters used are in Tables 1 and 2.

**Equivalence of stochastic and deterministic formalisms.** We first performed stochastic simulations with an initial latent cell pool size of $L_0$=$10^6$ cells, subjected to VRC01 therapy of a constant efficacy, $\varepsilon$=0.5, for a period of 21 days. The latent pool size is expected to lie in the range $10^5-10^8$ cells.[11,12] The low efficacy of $\varepsilon$=0.5 leads to rapid treatment failure, keeping the simulations tractable. We explore the implications of varying $\varepsilon$ below. Over the 21 d period studied, we found that the latent cell pool size, *L*, did not vary much from $L_0$ (Fig. S2(a)), consistent with experiments[13]. The waiting time for the first reactivation event, $\tau$, was small (Fig. S2(b)) and led to successful infection, characterized by a sustained rise in the productively infected cell population, *T*, in ~4 days (Fig. S2(c)), consistent with previous estimates in untreated individuals[14] given the low VRC01 efficacy assumed.

The formation of the first productively infected cell carrying a VRC01-resistant genome, $T_m$, had a waiting time probability density with a peak at ~7 d and a tail extending to 15 days (Fig. S2(d)). The corresponding probability density estimated using our deterministic formalism (

$\partial P(y < s)/\partial s = f_y(s)$) was in excellent agreement with stochastic simulations (Fig. S2(d)), giving us confidence in our simulations. As expected, increasing $L_0$ decreased the waiting times (Fig. S2(e)), increasing VRC01 efficacy increased the waiting times (Fig. S2(f)), and increasing viral fitness, achieved, for instance, with increasing viral production rate, decreased the waiting times (Fig. S2(g)). In all cases, the densities predicted by our deterministic calculations were in excellent agreement with the stochastic simulations. The agreement suggested that the deterministic model could be used to estimate the mean waiting time for VRC01-failure when the stochastic simulations became prohibitive.

**Estimates of VRC01 failure times.** With $L_0$~$10^7$ cells, representative of the latently infected pool size in chronically infected individuals[11,12], and with high VRC01 efficacies against the sensitive strains, $\varepsilon$ >0.95, based on the known $IC_{50}$ values and measured plasma bNAb concentrations[2], the waiting times were much larger than the 21 days period we simulated. We therefore used our deterministic framework to estimate the mean waiting time, $\tau_m$, for the formation of the first productively infected cell $T_m$. The model yielded $\tau_m$>100 d with $L_0$~$10^7$ cells and $\varepsilon$=0.98 (Fig. S2(h)), much larger than the VRC01 failure times observed clinically (2-4 weeks)[6]. $\tau_m$ did decrease with higher $L_0$ ($\geq 5\times10^7$ cells), but only a few people may be expected to have such large latent cell pools, whereas most individuals treated with VRC01 achieved failure within 2-4 weeks[6]. Further, the establishment of successful infection after the formation of the first $T_m$ may require crossing additional barriers, as mentioned above, so that $\tau_m$ is a lower bound on the failure time predicted by our model. We concluded, therefore, that VRC01 therapy failure is unlikely to be due to *de novo* mutation of wild-type strains contained in latently infected cells.

#### Supplementary Tables

**Table S1.** Relative replicative fitness, relative infectivity, and pre-existing frequencies of various mutants. Replicative fitness was estimated by analyzing competitive growth assays where available (Fig. S1) and by assuming zero epistasis otherwise. For the N279K and N280D mutants, competitive growth assays indicated replicative fitness indistinguishable from the wild-type. The relative infectivity of each strain was calculated as the ratio of its % infectivity to that of the wild type, reported elsewhere[2]. The frequencies were obtained using our model (Eqs. (1)-(10)).

| Strains | Replicative fitness | Infectivity | Frequency |
|---|---|---|---|
| YU2 (wild-type) | 1 | 1 | 0.999624 |
| N279K | 1 | 0.86 | $2.27\times10^{-4}$ |
| N280D | 1 | 0.453 | $5.8\times10^{-5}$ |
| R456W | 0.432 | 0.974 | $5.79\times10^{-5}$ |
| G458D | 0.491 | 0.1 | $3.3\times10^{-5}$ |
| N279K-N280D | 1 | 0.389 | $1.56\times10^{-9}$ |
| N279K-R456W | 1 | 0.427 | $1.59\times10^{-9}$ |
| N279K-G458D | 0.491 | 0.086 | $9.69\times10^{-10}$ |
| N280D-R456W | 0.432 | 0.441 | $1.16\times10^{-9}$ |
| N280D-G458D | 0.491 | 0.0453 | $9.48\times10^{-10}$ |
| R456W-G458D | 0.212 | 0.0974 | $9.56\times10^{-10}$ |
| N279K-N280D-R456W | 0.432 | 0.379 | $3.36\times10^{-14}$ |
| N279K-N280D-G458D | 0.491 | 0.0389 | $2.83\times10^{-14}$ |
| N279K-R456W-G458D | 0.491 | 0.0427 | $2.84\times10^{-14}$ |
| N280D-R456W-G458D | 0.212 | 0.0441 | $2.8\times10^{-14}$ |
| N279K-N280D-R456W-G458D | 0.433 | 0.208 | $9.2\times10^{-19}$ |

### Supplementary Figures

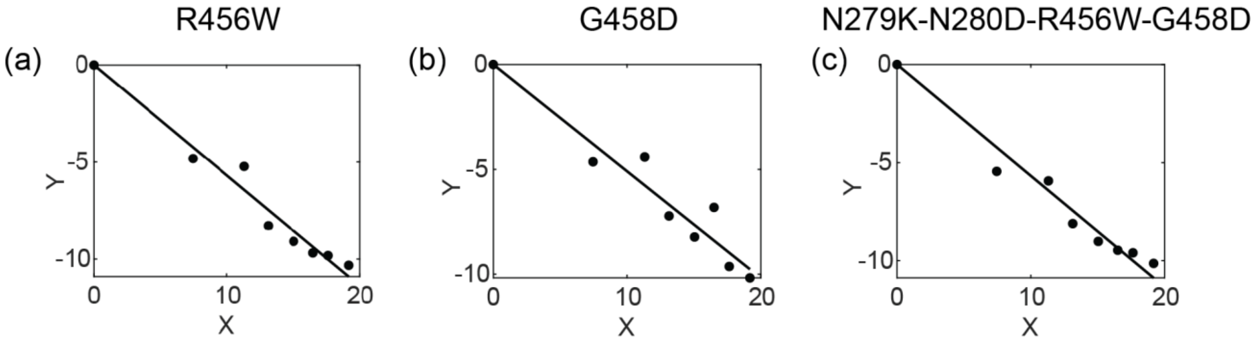


**Figure S1: Estimation of fitness using data from competitive growth assays.** We applied a previous formalism[15] to analyze competitive growth assays, where the selective advantage, *s*, of a mutant relative to the wild-type is given by $s = \ln\left[H(t)/H(0)\right] / \left[\ln\left[W(t)/W(0)\right] + \delta t\right]$, where $W(t)/W(0)$ is the fold expansion of wild type virus at time *t*; $H(t)/H(0)$ is the fold change of the mutant-to-wild type ratio at time *t*; and $\delta$ is the death rate of infected cells. Rewriting this equation as $Y = sX$, where $Y = \ln\left[H(t)/H(0)\right]$ and $X = \ln\left[W(t)/W(0)\right] + \delta t$, we estimate *s* by fits (lines) to corresponding data (symbols)[2] of *Y* vs. *X* for three mutants: **(a)** R456W, **(b)** G458D, **(c)** N279K-N280D-R456W-G458D. The best-fit estimates (95% CI) of *s* are -0.568 (-0.6089, -0.5272) for R456W; -0.5091 (-0.5662, -0.452) for G458D; and -0.5672 (-0.6075, -0.5269) for N279K-N280D-R456W-G458D. The relative replicative fitness of the respective strains are obtained as $\zeta = (1 + s)$.

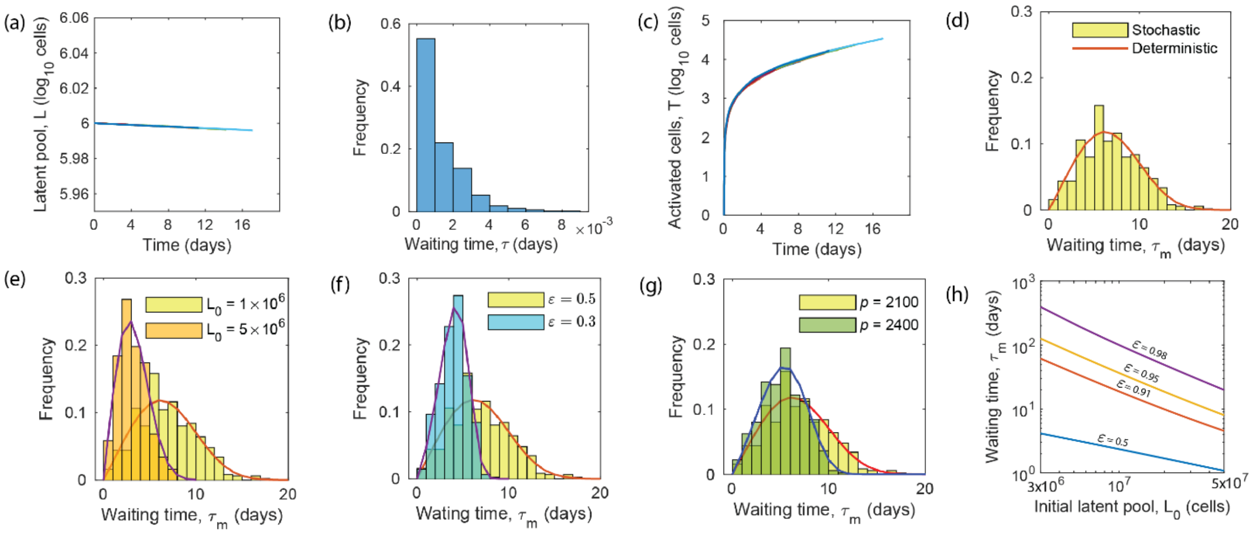


**Figure S2: VRC01 failure via *de novo* mutation.** Predictions of stochastic simulations (Eqs. S1-S11), showing **(a)** time-evolution of the latent cell pool, **(b)** distribution of the waiting time for the reactivation from latency, **(c)** time-evolution of activated or productively infected cells, and **(d)** distribution of the waiting time for the first productively infected cell carrying a VRC01-resistant strain (bar graph). (In (a) and (c), the different lines represent different stochastic realizations.) Variation of the latter distribution is shown with **(e)** initial latent cell pool size (cells), **(f)** VRC01 efficacy, and **(g)** viral production rate (virions/cell/day). In **(d)-(g)**, the corresponding probability density function calculated using the deterministic formalism (Eqs. S12-S17) is shown as solid lines. **(h)** Expected waiting time, $\tau_m$, for the formation of a productively infected cell carrying a VRC01-resistant provirus, calculated using the deterministic formalism as a function of the initial latent pool size for different values of the VRC01 efficacy indicated.

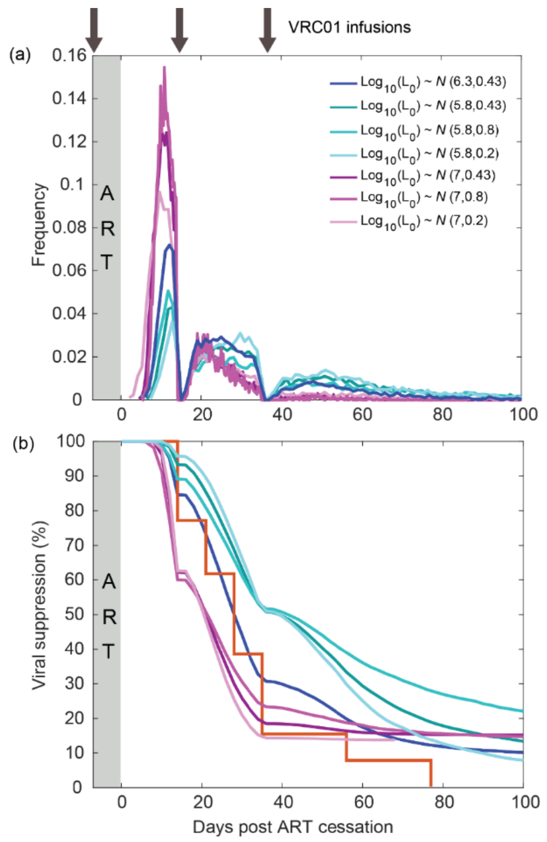


**Figure S3: Estimation of inter-patient variation in latent pool size.** Model predictions similar to those in **(a)** Fig. 4(c) and **(b)** Fig. 4(d) with virtual patient populations created by sampling $L_0$ (cells) from the different distributions indicated.  The red line in (b) is data from the A5340 trial.

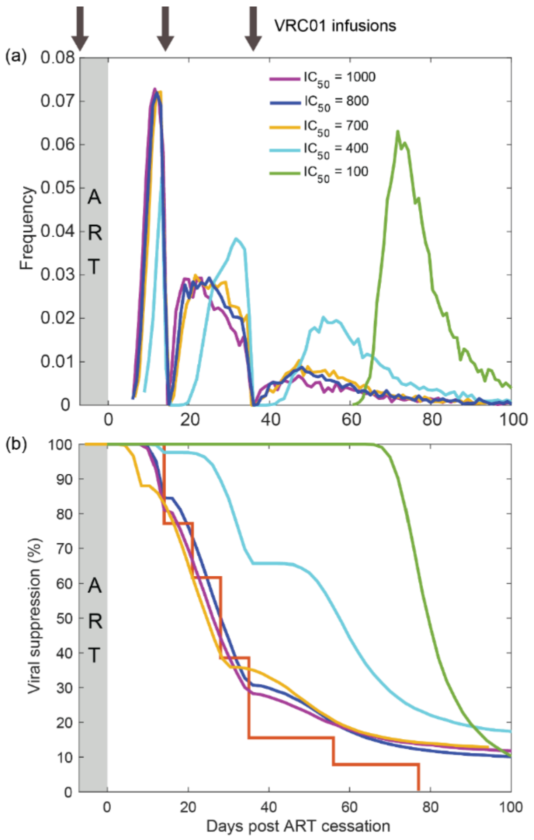


**Figure S4: Sensitivity to VRC01 *$IC_{50}$*.** Model predictions similar to those in **(a)** Fig. 4(c) and **(b)** Fig. 4(d) with different values of the $IC_{50}$ (µg/mL) indicated. The red line in (b) is data from the A5340 trial.

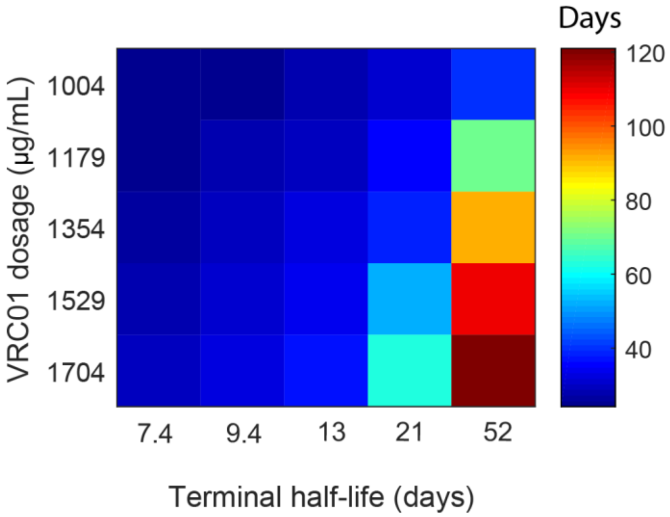


**Figure S5: Maximizing remission with VRC01**. Heat map showing the median rebound time, corresponding to the detection limit of 20 copies/mL, for different VRC01 dosages and half-lives. For each parameter combination, a virtual population of 10000 individuals was employed following the dosing schedule in the A5340 schedule, as in Fig. 4. Note that the modified bNAb VRC01LS has a >4-fold longer half-life than VRC01, and has been tested in healthy and uninfected adults for safety and pharmacokinetics[16]. The terminal half-life, $\eta_1$, has been varied accordingly. The initial total antibody concentration $(A_1 + A_2)$ has been varied while keeping the ratio $A_1 / A_2$ fixed, with the maximum $(A_1 + A_2)$ (in µg/mL) corresponding to the 40 mg/kg dosage of VRC01 (fitted from data[17]), which appears to be the maximum dosage of VRC01 reported. The other parameters are the same as in Fig. 4.

#### Supplementary references


1. Ramratnam, B. *et al.* Rapid production and clearance of HIV-1 and hepatitis C virus assessed by large volume plasma apheresis. *Lancet* **354**, 1782–1785 (1999).
2. Lynch, R. M. *et al.* HIV-1 Fitness cost associated with escape from the VRC01 class of CD4 binding site neutralizing antibodies. *J. Virol.* **89**, 4201–4213 (2015).
3. Hill, A. L., Rosenbloom, D. I. S., Fu, F., Nowak, M. A. & Siliciano, R. F. Predicting the outcomes of treatment to eradicate the latent reservoir for HIV-1. *Proc. Natl. Acad. Sci. U. S. A.* **111**, 13475–13480 (2014).
4. Conway, J. M., Perelson, A. S. & Li, J. Z. Predictions of time to HIV viral rebound following ART suspension that incorporate personal biomarkers. *PLoS Comput. Biol.* **15**, e1007229 (2019).
5. Gillespie, D. T. Exact stochastic simulation of coupled chemical reactions. *J. Phys. Chem.* **81**, 2340–2361 (1977).
6. Bar, K. J. *et al.* Effect of HIV antibody VRC01 on viral rebound after treatment interruption. *N. Engl. J. Med.* **375**, 2037–2050 (2016).
7. Arora, P. & Dixit, N. M. Timing the emergence of resistance to anti-HIV drugs with large genetic barriers. *PLoS Comput. Biol.* **5**, e1000305 (2009).
8. Nagaraja, P., Alexander, H. K., Bonhoeffer, S. & Dixit, N. M. Influence of recombination on acquisition and reversion of immune escape and compensatory mutations in HIV-1. *Epidemics* **14**, 11–25 (2016).
9. Perelson, A. S. & Ribeiro, R. M. Modeling the within-host dynamics of HIV infection. *BMC Biol.* **11**, 96 (2013).
10. Conway, J. M. & Coombs, D. A stochastic model of latently infected cell reactivation and viral blip generation in treated HIV patients. *PLoS Comput. Biol.* **7**, e1002033 (2011).
11. Chun, T. W. *et al.* Early establishment of a pool of latently infected, resting CD4+ T cells during primary HIV-1 infection. *Proc. Natl. Acad. Sci. U. S. A.* **95**, 8869–8873 (1998).
12. Estes, J. D. *et al.* Defining total-body AIDS-virus burden with implications for curative strategies. *Nat. Med.* **23**, 1271–1276 (2017).
13. Salantes, D. B. *et al.* HIV-1 latent reservoir size and diversity are stable following brief treatment interruption. *J. Clin. Invest.* **128**, 3102–3115 (2018).
14. Pinkevych, M. *et al.* HIV reactivation from latency after treatment interruption occurs on average every 5-8 days—implications for HIV remission. *PLOS Pathog.* **11**, e1005000 (2015).
15. Maree, A. F. M., Keulen, W., Boucher, C. A. B. & De Boer, R. J. Estimating relative fitness in viral competition experiments. *J. Virol.* **74**, 11067–11072 (2000).

16. Gaudinski, M. R. *et al.* Safety and pharmacokinetics of the Fc-modified HIV-1 human monoclonal antibody VRC01LS: A Phase 1 open-label clinical trial in healthy adults. *PLoS Med.* **15**, e1002493 (2018).

17. Lynch, R. M. *et al.* Virologic effects of broadly neutralizing antibody VRC01 administration during chronic HIV-1 infection. *Sci. Transl. Med.* **7**, 319ra206 (2015).
